# PARP and PI3K inhibitor combination therapy eradicates c-MYC-driven murine prostate cancers via cGAS/STING pathway activation within tumor-associated macrophages

**DOI:** 10.1101/2020.07.17.198598

**Authors:** Priyanka Dutta Gupta, Kiranj Chaudagar, Sweta Sharma-Saha, Kaela Bynoe, Lea Maillat, Brian Heiss, Walter M Stadler, Akash Patnaik

**Affiliations:** Section of Hematology/Oncology, Department of Medicine, University of Chicago, Chicago, IL; Translational and Clinical Research Institute, Newcastle University Centre for Cancer, Faculty of Medical Sciences, Newcastle upon Tyne, UK; Pritzker School of Molecular Engineering, University of Chicago, IL

**Keywords:** PARP inhibitor, PI3K inhibitor, c-MYC, STING, tumor-associated macrophages, immune checkpoint blockade

## Abstract

The majority of metastatic, castrate-resistant prostate cancer (mCRPC) patients are *de novo* resistant to immune checkpoint blockade (ICB), so therapeutic strategies to enhance immune-responsiveness are urgently needed. Here we performed a co-clinical trial of PARP inhibitor (PARPi) in combination with PD-1 or PDL-1 antibody in genomically unselected mCRPC patients or homologous-recombination proficient murine models, respectively, which demonstrated lack of efficacy. In contrast, PARPi in combination with PI3K inhibitor (PI3Ki), induced tumor regression via macrophage STING-dependent innate immune activation *in vivo*, and enhanced T-cell infiltration/activation in c-myc driven murine prostate cancer models, which was augmented by PD-L1 blockade. *Ex vivo* mechanistic studies revealed that PARPi-induced DNA double strand break-associated microvesicles released from tumor cells, coupled with PI3Ki-mediated c-GAS de-repression, were both required for macrophage cGAS/STING pathway activation. These data demonstrate that PARPi/PI3Ki combination triggers macrophage STING-mediated anti-cancer innate immunity, which is sufficient to induce tumor regression in ICB-refractory c-myc-driven prostate cancer.

**STATEMENT OF SIGNIFICANCE:** Co-targeting of PARP and PI3K signaling pathways activates c-GAS/STING pathway within tumor-associated macrophages, thereby enhancing T cell recruitment/activation and cancer clearance in c-myc-driven murine prostate cancer models. PARPi/PI3Ki combination therapy could markedly increase the fraction of mCRPC patients responsive to ICB, independent of germline or tumor homologous recombination status.

## INTRODUCTION

Prostate Cancer (PC) is the most common malignant neoplasm in men and the second most frequent cause of cancer death for males in the United States. While there have been incremental advances, mCRPC remains an incurable disease with high morbidity and mortality (1, 2), so there is an urgent need to develop definitive treatments that improve survival. Over the past decade, there has been a resurgent interest in cancer immunotherapy, partly based on the profound and durable clinical responses to immune checkpoint blockade (ICB) antibodies targeting CTLA-4 and PD-1/PD-L1 (3). However, only approximately 10-25% of mCRPC patients respond to these approaches (4–6).

*MYC* is a “master” proto-oncogene that contributes to tumorigenesis of greater than 75% of all advanced refractory human cancers, particularly prostate, colon, breast, cervical cancers, acute myeloid leukemia, lymphomas, small-cell lung cancer, and neuroblastoma (7, 8). C-myc is a transcription factor encoded by the *MYC* gene on locus 8q24.21, which is frequently amplified in human cancers (8). However despite multiple pharmaceutical efforts, c-myc has remained “undruggable” (9–11). Furthermore, c-myc driven-cancers are resistant to ICB (12). Therefore, therapeutic strategies that target c-myc-driven cancers and enhance their responsiveness to ICB are urgently needed.

The cGAS/STING pathway is known to be physiologically activated by cytosolic double-stranded DNA (dsDNA), which typically occurs in the context of viral infections, resulting in the generation of cytosolic cyclic dinucleotides generated by the Cyclic GMP-AMP synthase (cGAS) enzyme, downstream activation of the Stimulator of Interferon Genes (STING) pathway and induction of Type I interferon (IFN) production (13–15). Recent preclinical studies have demonstrated that PARPi, which are FDA approved for BRCA1/2 mutated prostate (16, 17), breast (18), ovarian (19) and pancreatic cancers (20), can activate the innate immune cGAS/STING pathway in murine homologous recombination (HR)- deficient breast and ovarian cancer models, resulting in sensitization of these tumors to ICB (21, 22). Furthermore, preclinical data suggests that PARPi can elicit DNA damage in HR-proficient cancers (23), but this is generally insufficient for meaningful clinical activity (24).

To test the hypothesis that PARPi and resulting DNA damage can sensitize HR-proficient mCRPC to ICB, we performed a co-clinical trial testing the combination of PARPi with PD-1 or PD-L1 antibody, in both HR-proficient mCRPC patients and murine models, respectively, which demonstrated lack of efficacy. In contrast, concomitant PI3K inhibitor (PI3Ki) treatment with PARPi induced tumor regression in c-myc driven murine PC models, via tumor cell-extrinsic, macrophage STING-dependent innate immune activation, which was accompanied by enhanced T-cell infiltration/activation. Critically, the anti-tumor response elicited by PARPi/PI3Ki was augmented by PD-1/PDL1 axis blockade and abrogated in immunodeficient mice and immunocompetent mice treated with systemic macrophage depleting agent (Clodronate) or STING antagonist (H-151). Mechanistically, we observed that DNA double-strand break (DSB)-associated MVs released from PARPi-treated transgenic c-myc over-expressing cancer cells, along with concomitant PI3Ki-mediated post-translational de-repression of cGAS enzymatic activity, increased cGAMP and activated the STING pathway within tumor-associated macrophages (TAMs). Taken together, these data demonstrate that PARPi/PI3Ki combination drives anti-cancer innate immunity via cGAS/STING pathway activation within TAMs, resulting in tumor regression in murine models of c-myc driven PC. Based on these observations, clinical trials testing PARPi/PI3K inhibitors with ICB are warranted in immunotherapy-refractory HR-proficient advanced PC.

## RESULTS

### The sparse immune infiltrates in human mCRPC and murine myc-driven PC models are dominated by myeloid suppressive cells, particularly tumor-associated macrophages (TAMs)

As a first step towards deconvoluting the complex ecosystem of the metastatic tumor immune microenvironment in mCRPC, we performed flow cytometric analysis of 4 tumors isolated from human mCRPC lymph node biopsy samples. Notably, immune profiling revealed an “immune desert” with a paucity of CD45+ cells within the TME **(Fig. 1A, Supplementary Table 1)**. Furthermore, the small fraction of immune cells within the TME were predominantly composed of CD11b+ myeloid cells (approx. 80%, **Fig. 1B**), with F4/80+ TAMs comprising the highest frequency of CD45+ cells **(Fig. 1C)**. Approximately 80% of the F4/80+ TAMs within the mCRPC samples were HLA-DR^-^/MHC II^-^ **(Fig. 1D)**, indicating that these cells are unactivated/immunosuppressive M2-like macrophages. We additionally performed immune profiling of tumors derived from two transgenic c-myc-driven prostate cancer lines, Myc-CAP (25) and B6-Myc (26), generated in FVB/NJ and C57Bl/6J genetic backgrounds, respectively. We observed a similar paucity of CD45+ immune cells within the TME, which was approx. 10-fold lower (p<0.001) in Myc-CAP **(Fig. 1E)**, relative to the B6-Myc tumors **(Fig. 1I)**. However, similar to mCRPC samples, both Myc-CAP and B6-Myc tumors showed a relative predominance (approx. 80% of CD45+ immune cells) of CD11b+ myeloid cells **(Fig. 1B, 1F, 1J)** within the TME. Within the CD11b+ myeloid population, F4/80+ TAMs comprise the predominant immune subset **(Fig. 1G, 1K),** and 65-70% of these cells were unactivated/immunosuppressive (M2) HLA-DR^-^/MHC II^-^ macrophages **(Fig. 1H, 1L),** similar to what was observed in the mCRPC patient samples **(Fig. 1D)**. Taken together, these data demonstrate similar immune contexture in human mCRPC and murine c-myc-driven cancers, which is dominated by myeloid suppressive cells, particularly TAMs.

**Figure 1:**
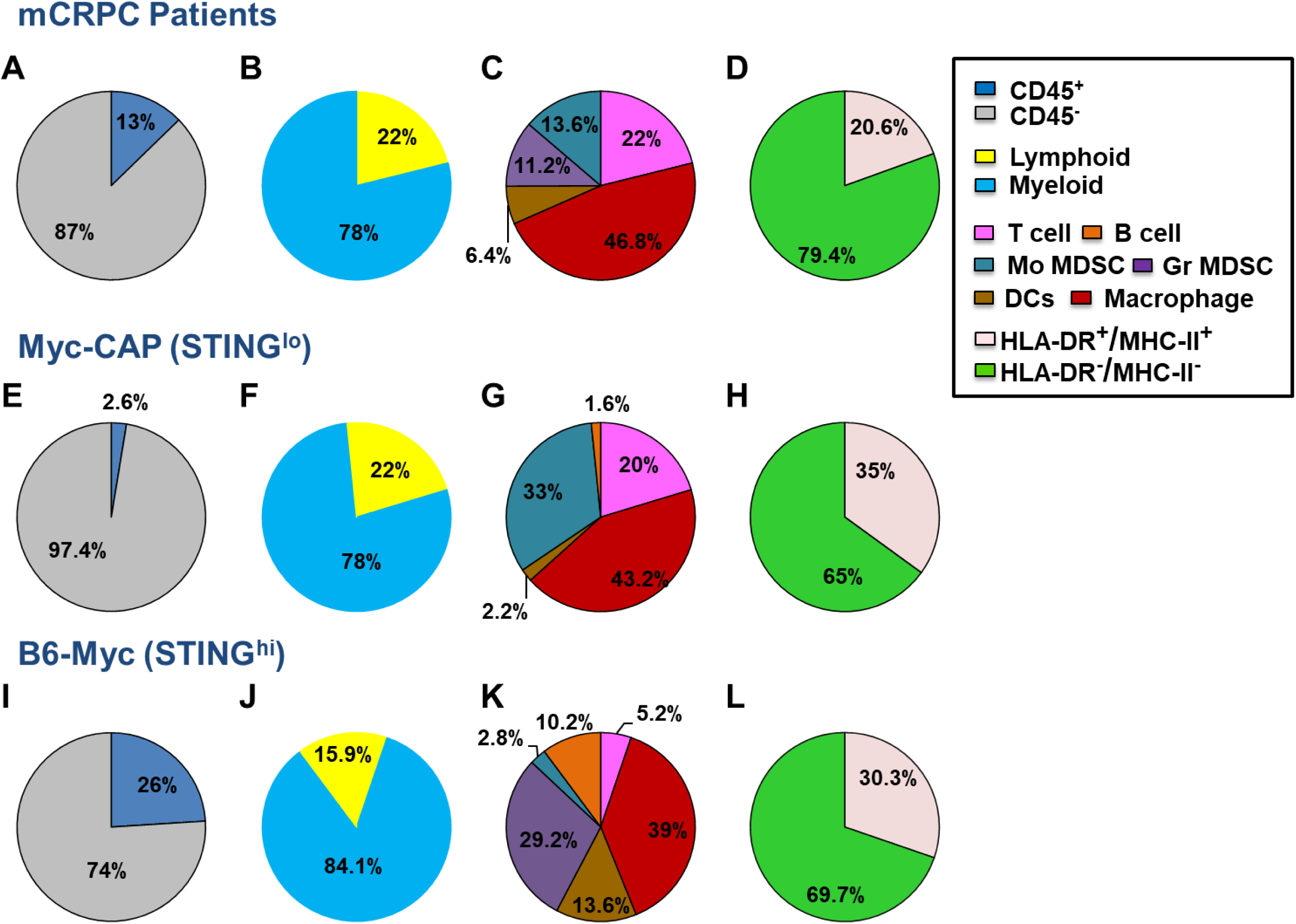
The immune infiltrates within human mCRPC and murine c-myc-driven PC tumors are dominated by tumor-associated macrophages (TAMs). Single cell suspensions from human mCRPC biopsies, murine Myc-CAP and B6-Myc syngeneic tumors were stained with anti-human/mouse lineage-specific antibodies and analyzed by flow cytometry. (A,E,I) Tumor cells (CD45-) vs. Immune cells (CD45+); (B,F,J) Lymphoid - T (CD3+) + B (CD19+) cells vs. Myeloid (CD11b+; macrophages, DCs, MDSCs – monocytic and granulocytic) cells. (C,G,K) Lymphoid and Myeloid compartments were analyzed for individual immune cell subsets using the following additional markers: Macrophages (hCD45+CD11b+CD163+CD68+/mCD45+CD11b+F480+), DCs (hCD45+CD11b+ CD11c+/mCD45+CD11b+CD11c+), Gr-MDSCs (hCD45+CD11b+HLA-DR-CD15^hi^ CD33^lo^/mCD45+CD11b+MHC-II-Ly6G^hi^Ly6C^lo^), Mo-MDSCs (hCD45+CD11b+HLA-DR-CD15^lo^CD33^hi^/mCD45+CD11b+MHC-II-Ly6G^lo^Ly6C^hi^); (D,H,L) HLA-DR expression on TAMs in mCRPC & MHC-II expression on TAMs within TME of murine tumors were analyzed; n=4 for human and n= 3-4 for murine samples. DC=dendritic cells; MDSC= myeloid derived suppressor cell, h= human, m= mouse.

### Expression of STING and an activated myeloid gene expression within primary PC samples positively correlate with biochemical recurrence-free survival

We next assessed cGAS and STING expression in primary human PC samples within the TCGA, and discovered reduced expression relative to normal tissue counterparts **(Fig. 2A)**. Furthermore, transcriptomic data analysis of primary PC samples within the TCGA, revealed that high risk (Gleason ≥8) patients with higher biochemical recurrence **(Fig. 2B),** had decreased gene expression of STING **(Fig. 2C)**, myeloid activation markers HLA-DR and CD86, (**Fig. 2D-E**), and T cell chemotactic factors CXCL10 and CCL5 (**Fig. 2F-G)**, relative to low-risk (Gleason 6/7) patients. These data identify a positive correlation between STING expression, myeloid activation states, T cell chemotactic factor expression, and clinical outcome.

**Figure 2:**
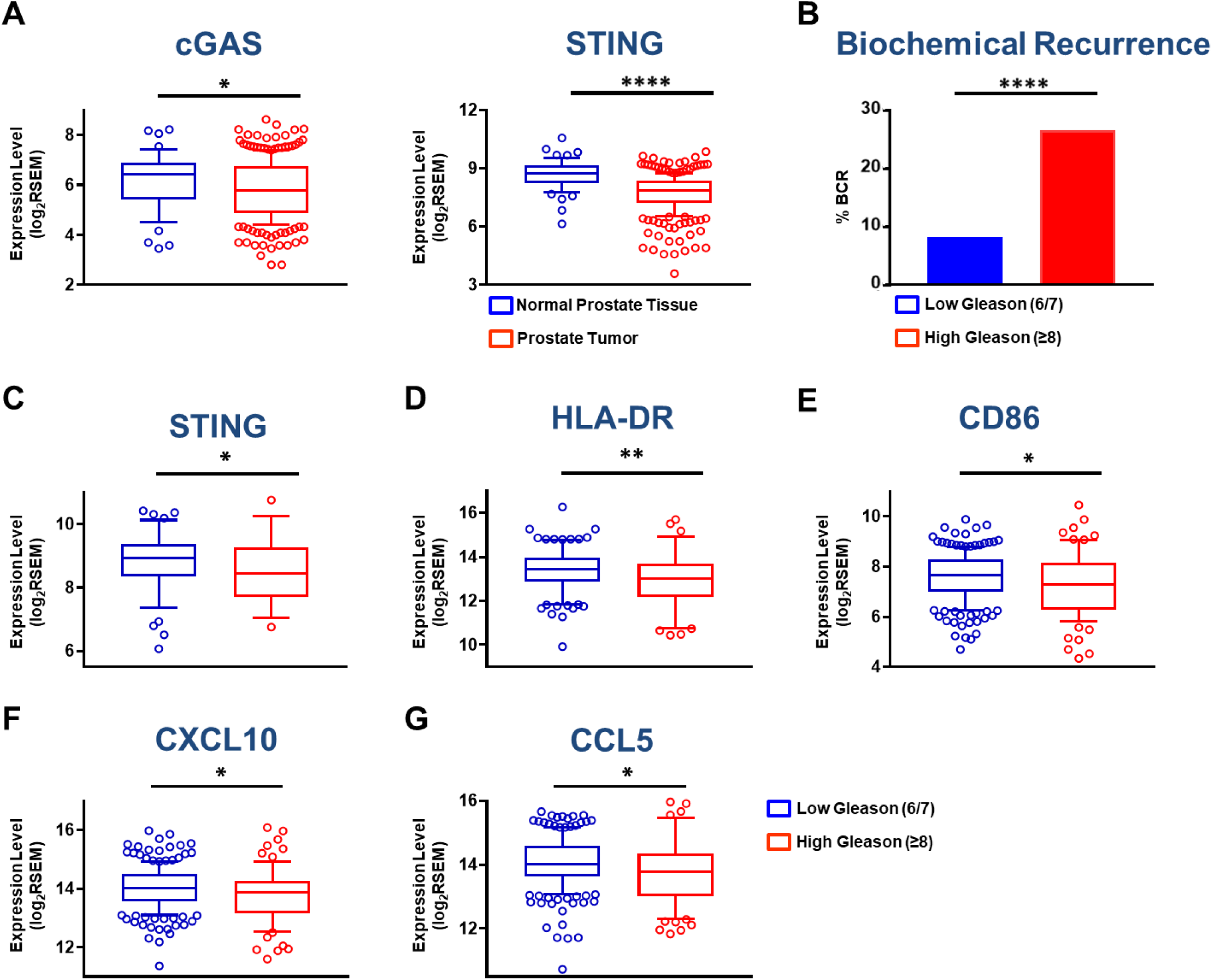
High-risk PC patients exhibit reduced STING, myeloid activation and T cell chemotactic factor gene expression. **(A)** cGAS and STING expression in normal vs. PC within the TCGA. **(B)** PC patients in TCGA were further subdivided into Low and High Gleason score groups, defined as follows: Low Gleason score = 6/7 and High Gleason Score ≥8. Frequency of biochemical recurrence (BCR) for evaluable cases within these cohorts was determined and differential gene expression across the two groups was further analyzed for STING **(C)**, indicated myeloid activation-specific genes **(D-E)** and T cell chemotactic factors **(F-G)** secreted downstream of STING activation; n= 290 (Low Gleason group n= 211; High Gleason group n= 79); Significance/p-values were calculated using Mann Whitney/Un-paired t-test for panels (A, C-G) and Kolmogorov Smirnov test for panel (B) and indicated as follows, *p < 0.05; **p < 0.01; ****p<0.0001. BCR=BioChemical Recurrence.

Next we assessed cGAS and STING protein expression in Myc-CAP and B6-Myc cells, and observed low and high cGAS/STING expression in Myc-CAP (STING^lo^) and B6-Myc (STING^hi^) cells, respectively **(Supplementary Fig. 1A)**, which mimic the STING gene expression patterns observed in low and high-risk PC subgroups groups within the TCGA (Fig. 2). Consistent with the relative expression data of STING pathway components in B6-Myc (STING^hi^) cancer cells, we observed that deliberate STING activation with DMXAA (mouse STING agonist) activated the cGAS/STING signaling pathway components phospho-TANK-binding kinase 1 (p-TBK1) and phospho-interferon regulatory factor-3 (p-IRF3) in B6-Myc (STING^hi^) cells **(Supplementary Fig. 1A)**, not observed in Myc-CAP (STING^lo^) cells. DMXAA treatment of bone marrow derived macrophages (BMDMs) also activated the c-GAS/STING signaling pathway **(Supplementary Fig. 1B)**. Furthermore, ELISA analysis revealed that DMXAA treatment of B6-Myc (STING^hi^) cancer cells and BMDMs elicited a 48.1-fold and 46.4-fold induction in IFN-β levels, not observed in Myc-CAP (STING^lo^) cancer cells **(Supplementary Fig. 1C, 1D).** Interestingly, *ex vivo* treatment of single cell suspensions derived from Myc-CAP (STING^lo^) tumors with DMXAA demonstrated a 11.4-fold increase in IFN-β levels within the TME, but a lack of response specifically within the CD45-negative tumor cell fraction **(Supplementary Fig 1E)**. Collectively, these data demonstrate that STING pathway can be activated in a tumor cell *extrinsic* manner within Myc-CAP (STING^lo^) tumors. On the other hand, B6-Myc (STING^hi^) tumors can turn on the STING pathway in both tumor cell intrinsic and extrinsic compartments.

### Concomitant PI3Ki sensitizes B6-Myc (STING^hi^) murine PC to PARPi/aPD-L1 combination therapy, via STING-dependent, TAM-driven immune mechanism

Recent preclinical data has demonstrated that PARP inhibitor (PARPi)- induced DNA damage can reprogram the tumor immune microenvironment via tumor cell intrinsic cGAS/STING activation, thereby enhancing T cell infiltration and efficacy of ICB in homologous recombination deficient (HRD) breast (21) and ovarian (22) cancer models. Given preclinical data suggesting that PARPi can induce DNA damage in HR-proficient cancers (23), we hypothesized that PARPi-induced DNA damage can enhance ICB efficacy, independent of HR status, in mCRPC patients enrolled in an investigator-initiated co-clinical trial at the University of Chicago (NCT03572478, IRB18-0154). To test this hypothesis, mCRPC patients that had progressed on at least one-line of AR-targeted therapy in the castrate-resistant setting, were treated with PARPi rucaparib (Clovis Oncology) in combination with nivolumab (Bristol Myers Squibb) until disease progression and/or unacceptable toxicity. A Waterfall plot for mCRPC patients on study for at least 90 days, demonstrated that only 1 of 7 evaluable patients responded to the combination therapy **(Fig. 3A)**. The single responder patient harbored a BRCA2 mutation that was predicted to respond to PARPi monotherapy, while the remaining patients had an HRD-proficient tumor mutational status. The combination of rucaparib and nivolumab had a PSA response rate of 9% (1/11 patients) and an objective response rate, per RECIST/PCWG3 criteria, of 0% (0/11 patients). Median progression-free survival for mCRPC patients on trial was 2.96 months (95% Confidence Interval, 2.03 months-not assessable). Taken together, these data demonstrate that the majority of patients did not exhibit clinically meaningful responses to rucaparib/nivolumab combination therapy. Consistent with our clinical trial data, we observed *de novo* resistance of B6-Myc (STING^hi^) syngeneic tumors to rucaparib or rucaparib/PD-L1 antibody combination **(Fig. 3B)**.

**Figure 3:**
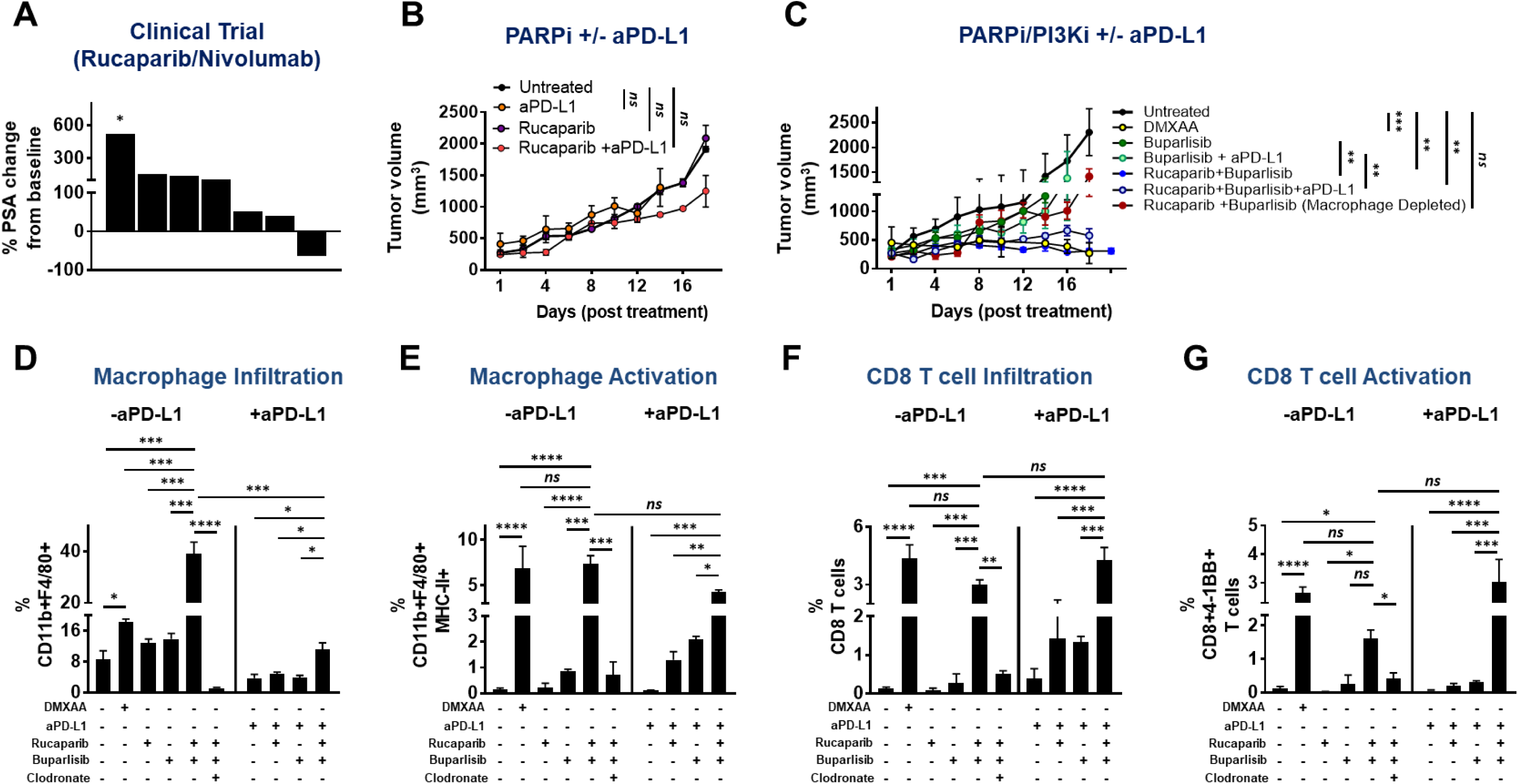
PARPi/PD-1 targeted combination therapy shows lack of efficacy in PC co-clinical trials, which can be reversed by concomitant PI3Ki treatment. **(A)** mCRPC patients were enrolled in a Phase Ib rucaparib/nivolumab co-clinical trial at University of Chicago, and treated until disease progression or unacceptable toxicity. As part of the study, blood was processed to collect sera for ELISA based determination of PSA levels at baseline and every month on study. Data obtained was used to calculate percent change in PSA levels at approximately 90 days, relative to baseline values. Each bar represents a single patient. * indicates patient who progressed early at 41 days, as per RECIST guidelines; n = 7 patients. **(B-C)** B6-Myc (STING^hi^) tumor-bearing syngeneic mice were treated with the indicated drug(s) and euthanized when untreated tumors reached approx. 2500 mm^3^. Tumor volume curves for duration of treatment are shown. **(D-G)** Single-cell suspensions were generated from harvested tumors and analyzed by flow cytometry for the indicated immune cell populations and their activation states. Data are represented relative to CD45+ immune cells. n=3 animals per treatment group from 2 independent experiments. Significance/p-values were calculated by one-way ANOVA/paired t-test and indicated as follows: *p < 0.05; **p < 0.01; ***p < 0.001; ns = not statistically significant.

To address the mechanistic basis for why PARPi was insufficient to sensitize B6-Myc (STING^hi^) tumors, we treated B6-Myc *in vitro* with rucaparib at 500 nM concentration that completely inhibits PARylation, and evaluated its impact on DNA damage, as assessed by quantification of p-γH2AX foci (marker of dsDNA breaks) using confocal microscopy. Interestingly, B6-Myc cells did not show a statistically significant increase in DNA double-strand breaks (DSBs) following single-agent rucaparib treatment **(Supplementary Fig. 2A).** Several studies have demonstrated that combination of PARP inhibitors (PARPi) and pan-PI3K (PI3Ki) inhibitors induce additive DNA damage and tumor regression in prostate and endometrial cancers (27, 28). Consistent with these prior observations, we observed a statistically significant additive increase in DNA DSBs following concomitant treatment with rucaparib and buparlisib (pan-PI3K inhibitor), but not with corresponding single-agent treatments in B6-Myc (STING^hi^) cells *in vitro* **(Supplementary Fig. 2A)**. Using concentrations for rucaparib and buparlisib of 500 nM and 1 μM, respectively, that achieved complete target inhibition *in vitro* (**Supplementary Fig. 2B)**, drug combination studies revealed that there was no change in viability of B6-Myc cells **(Supplementary Fig. 2C)**. Next we tested the impact of rucaparib and buparlisib combination, with or without PD-L1 antibody, on the ability to control tumor growth of B6-Myc (STING^hi^) syngeneic mice *in vivo*. At doses that pharmacodynamically inhibit PARP and PI3K enzymatic activity within the tumor *in vivo* **(Supplementary Fig. 3A)**, we observed complete tumor regression with the rucaparib/buparlisib combination relative to either single-agent, that was maintained with the addition of PD-L1 antibody. This tumor clearance and immune activation elicited by the rucaparib/buparlisib combination was phenocopied by DMXAA (mouse STING agonist) administration **(Fig. 3C-G, Supplementary Fig 6A and 6B)**. To address the discordance between *in vitro* cytotoxicity data and *in vivo* anti-cancer responses observed with rucaparib/buparlisib combination, we hypothesized that rucaparib/buparlisib combination is working predominantly via a tumor cell extrinsic immune mechanism. To test this possibility, we evaluated the impact of the rucaparib/buparlisib combination therapy in immunodeficient athymic nude mice implanted with B6-Myc (STING^hi^) allograft tumors. Strikingly, the anti-cancer mechanism of rucaparib/buparlisib was abolished in immunodeficient athymic nude mice **(Supplementary Fig. 4)**, thus demonstrating that this combination drives tumor regression via a cancer cell non-autonomous immune mechanism.

Immune TME profiling studies revealed that rucaparib/buparlisib combination in syngeneic B6-Myc (STING^hi^) demonstrated an increase in macrophage infiltration **(Fig. 3D)** and activation **(Fig. 3E)**, but not dendritic cell (DC) activation **(Supplementary Fig. 5)** and was accompanied by an increase in CD4 and CD8 T cell infiltration **(Supplementary Fig. 6A, Fig. 3F)** and activation **(Supplementary Fig. 6B, Fig. 3G)**, respectively, relative to corresponding single-agent controls. Furthermore, the addition of PD-L1 antibody to rucaparib/buparlisib combination accentuated the anti-tumor immune responses, particularly CD4 infiltration, relative to buparlisib/rucaparib treatment **(Supplementary Fig. 6A)**. Critically, the immunologic changes within the TME and tumor clearance elicited by PARPi/PI3Ki treatment were significantly attenuated by systemic macrophage depletion with clodronate **(Fig. 3C-G, Supplementary Fig 6A-B),** suggesting a macrophage-driven (DC-independent) anti-cancer innate immune mechanism for this combination. Furthermore, the anti-tumor immune response elicited by rucaparib/buparlisib combination increased MHC Class I expression within tumor cells, which was also suppressed by clodronate **(Supplementary Fig. 7),** likely related to suppression of IFN-ϒ-producing T cell infiltration/activation following TAM depletion.

To elucidate the role of STING pathway activation and the relative contributions of tumor cell intrinsic vs. extrinsic STING on the observed tumor regression, B6-Myc (STING^hi^) tumor allografts were implanted into STING^-/-^ C57Bl/6J mice. Strikingly, PARPi/PI3Ki-mediated B6-Myc tumor regression was partially attenuated in STING^-/-^ C57BL6 mice, and resulted in reduced macrophage and T cell activation, relative to their STING^+/+^ counterparts **(Supplementary Fig. 8)**. Taken together, these results demonstrate that PARPi/PI3Ki can induce tumor regression in B6-Myc (STING^hi^) syngeneic model via an innate immune mechanism that is driven by tumor cell extrinsic host STING within TAMs.

### PARPi/PI3Ki in combination with androgen deprivation therapy (ADT) causes tumor regression *in vivo* in Myc-CAP (STING^lo^) tumors, which is driven by TAM-mediated anti-cancer innate immunity

While PARPi/PI3Ki was sufficient to induce tumor clearance in B6-Myc (STING^hi^) syngeneic mice, Myc-CAP (STING^lo^) syngeneic mice are *de novo* resistant to this combination **(Supplementary Fig. 9)**. This could be related to the growth of these tumors in the FVB/NJ genetic background, which is known to have low immunogenicity (29–31). Prior studies have demonstrated that early stages of castration, which is standard-of-care for advanced prostate cancer (2), induce T-effector and T-regulatory cell infiltration within prostate tumors (32, 33). Consistent with prior data, we observed an approximately 5-fold increase in CD45+ infiltration in Myc-CAP (STING^lo^) tumors following castration, predominantly driven by an increase in the TAM subset, with a smaller contribution of CD4+ T cell subsets, but not CD8+ T cells **(Supplementary Fig. 10A-D)**. In addition, there is an increased PD-L1 expression in CD45-fractions and CD45+ (particularly TAMs) within the TME following castration **(Supplementary Fig. 10E-G)**. However, castration alone or in combination with rucaparib and/or PD-L1 antibody was insufficient to control tumor growth in Myc-CAP (STING^lo^) syngeneic mice **(Fig. 4A)**, which was consistent with our recent co-clinical trial data of rucaparib/nivolumab in mCRPC patients **(Fig. 3A)**.

**Figure 4:**
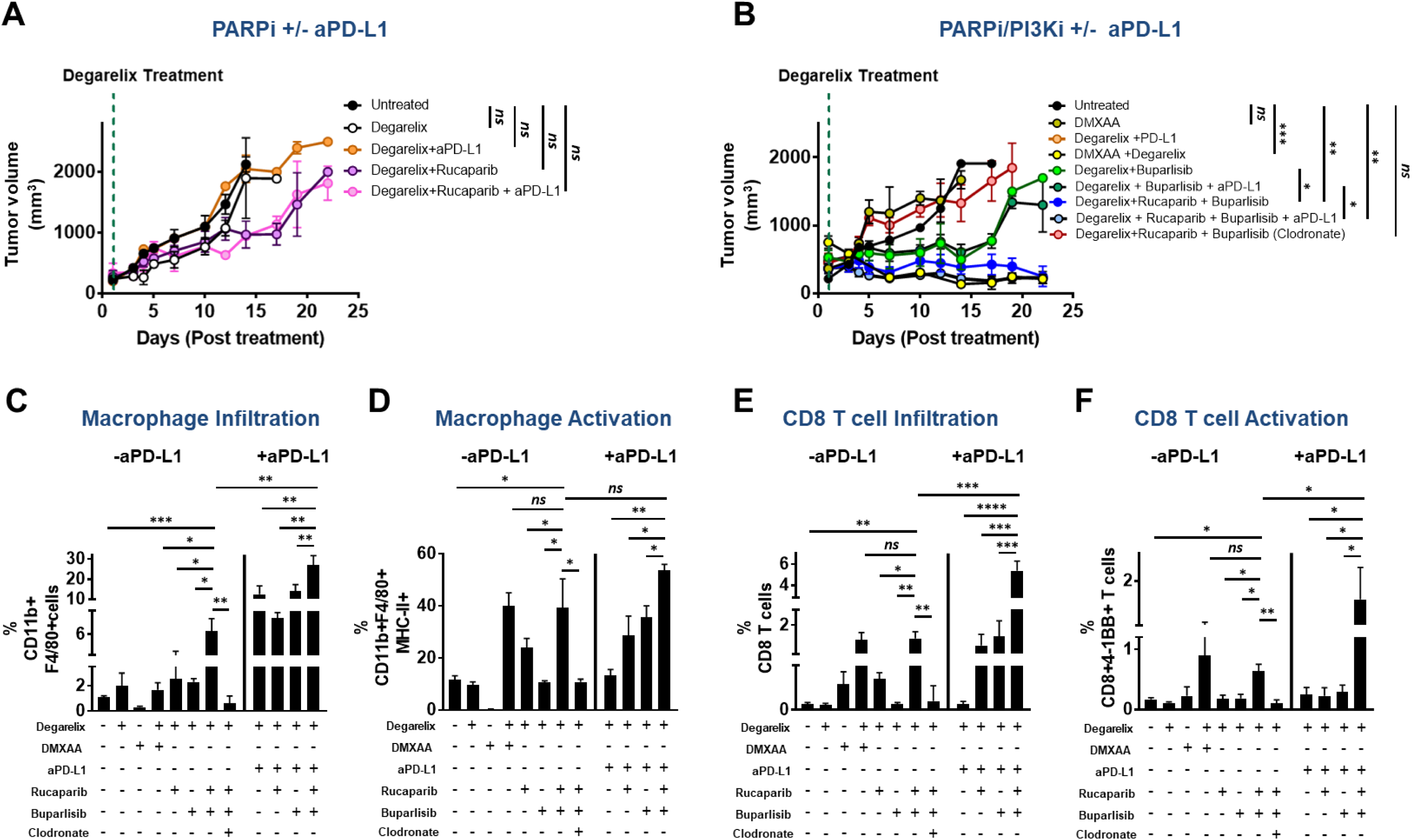
ADT/PARPi/PI3Ki combination induces TAM-driven tumor control in Myc-CAP (STING^lo^) syngeneic mice. **(A-B)** Myc-CAP (STING^lo^) tumor-bearing syngeneic mice were treated with the indicated drug(s) until untreated tumors reached approx. 2500 mm^3^. Tumor volume curves for duration of treatment are shown. **(C-F)** Single-cell suspensions were generated from harvested tumors and analyzed by flow cytometry for the indicated immune cell populations and their activation states. Data are represented relative to CD45+ immune cells. n=3-5 animals per treatment group from 2 independent experiments. Significance/p-values were calculated by one-way ANOVA and are indicated as follows: *p < 0.05; **p < 0.01, ***p < 0.001; ****p<0.0001; ns= not statistically significant.

We next tested the hypothesis that the immunostimulatory effects of castration would lower the threshold for the PARPi/PI3Ki combination to elicit a potent anti-tumor response in Myc-CAP (STING^lo^) syngeneic mice. Strikingly, we observed that the combination of degarelix and rucaparib/buparlisib resulted in complete tumor regression in Myc-CAP (STING^lo^) syngeneic mice, not observed with single agent degarelix, rucaparib and buparlisib or their corresponding doublet therapies **(Fig. 4B)**. The combination of degarelix and DMXAA phenocopied the complete tumor clearance observed with degarelix/rucaparib/buparlisib. Importantly, the anti-tumor response observed with degarelix/rucaparib/buparlisib was abolished in immunodeficient athymic nude **(Supplementary Fig 11A)** and NOD/SCID mice **(Supplementary Fig. 11B)**, thus demonstrating that this regimen induces tumor control via an immune-dependent mechanism. Furthermore, degarelix/rucaparib/buparlisib triple combination led to an increase in macrophage infiltration **(Fig. 4C)**, and activation **(Fig. 4D),** which was accompanied by an increase in CD4 and CD8 T cell infiltration **(Supplementary Fig. 12A, Fig. 4E)** and activation **(Supplementary Fig. 12B, Fig. 4F)**, respectively, relative to corresponding singlet or doublet controls. Furthermore, this immune activation effect of degarelix/rucaparib/buparlisib was accentuated by PD-L1 antibody, specifically with respect to macrophage infiltration and CD8 infiltration/activation **(Fig 4C, 4E, 4F)**. Gene expression (qRT-PCR) analysis and flow cytometric analysis of tumor extracts revealed increased *il12b* expression **(Supplementary Fig. 13A)**, and decreased Arginase-I expression **(Supplementary Fig. 13B)**, respectively, within TAMs from degarelix/rucaparib/buparlisib-treated tumors, relative to corresponding singlet or doublet controls, thus demonstrating that the triple combination enhanced M1 macrophage polarization within the TME. Furthermore, we observed that concomitant clodronate treatment abolished the tumor regression and immune-permissive reprogramming observed with degarelix/rucaparib/buparlisib treatment, with or without PD-L1 antibody **(Fig. 4B-F, Supplementary Fig. 12A-B**), similar to what was observed in B6-Myc (STING^hi^) model. Taken together, these data demonstrate that ADT/PARPi/PI3Ki combination induces a macrophage-mediated innate immune response in Myc-CAP (STING^lo^) syngeneic model, resulting in tumor clearance. Since the ADT/PARPi/PI3Ki-mediated tumor clearance was abolished in athymic nude mice, the anti-cancer responses were mediated at least in part by macrophage-mediated activation of T cell immunity.

### PARPi/PI3Ki-induced STING Pathway Activation Within TAMs is Mediated via MVs Released from Tumor cells

To determine whether PARPi/PI3Ki combination elicits DNA DSBs in Myc-CAP (STING^lo^) context, cells were treated *in vitro* with PARPi rucaparib, singly and in combination with buparlisib, at their respective target inhibitory concentrations **(Supplementary Fig. 14)**. Interestingly, we observed that treatment of Myc-CAP cells with rucaparib caused an increase in DNA DSBs, as measured by number of p-γH2AX foci, which was not enhanced by the addition of PI3Ki **(Supplementary Fig. 14A**). In addition, there was no change in the viability of Myc-CAP cells treated at complete target inhibition concentrations of 500 nM and 1 μM for rucaparib and buparlisib (**Supplementary Fig. 14B)** respectively, singly and in combination **(Supplementary Fig. 14C)**. Given the requirement of dual PARP and PI3K inhibition for tumor regression *in vivo*, these *in vitro* data suggest that concomitant PI3Ki treatment activates non-tumor cell autonomous, TAM-mediated innate immunity via DNA-damage independent mechanism in Myc-CAP (STING^lo^) tumors.

To elucidate the mechanism by which PARPi/PI3Ki induces STING pathway activation within TAMs, we tested the hypothesis that rucaparib/buparlisib-induced DNA DSBs within tumor cells cross-talks via MVs that activate cGAS/STING pathway within TAMs in the TME. We first evaluated the quantity and cargo content of MVs isolated from B6-Myc (STING^hi^) and Myc-CAP (STING^lo^) cells treated with rucaparib, singly and in combination with buparlisib. We observed that the MVs ranged from 50-100 nm in size, based on Nanoparticle Tracking Analysis (NTA) measurements. Futhermore, the quantity of DNA DSBs associated with MVs, was directly proportional to intracellular DNA damage content, as assessed by Nanodrop and p-ϒH2AX foci quantification respectively, for both B6-Myc (STING^hi^) and Myc-CAP (STING^lo^) cells treated with rucaparib and buparlisib, singly and in combination **(Supplementary Fig. 2A, 2D 14A, 14D)**.

Next we conducted *ex vivo* assays using single-cell suspensions of Myc-CAP (STING^lo^) tumors treated with exogenous rucaparib, singly and in combination with buparlisib. Interestingly, we observed an increase in IFNβ production within the supernatants of tumor allograft single cell suspensions following *ex vivo* rucaparib/buparlisib combination treatment, not observed with either single agent. Furthermore, we tested the impact of these drug(s) in the presence or absence of GW4869, an inhibitor of MV biogenesis and release. Concomitant *ex vivo* GW4869 treatment of tumor cell suspensions abolished IFNβ production following rucaparib/buparlisib treatment **(Fig. 5A)**, thus demonstrating that cGAS/STING pathway activation within the TME occurs via MV-associated DNA DSBs released from tumor cells.

**Figure 5:**
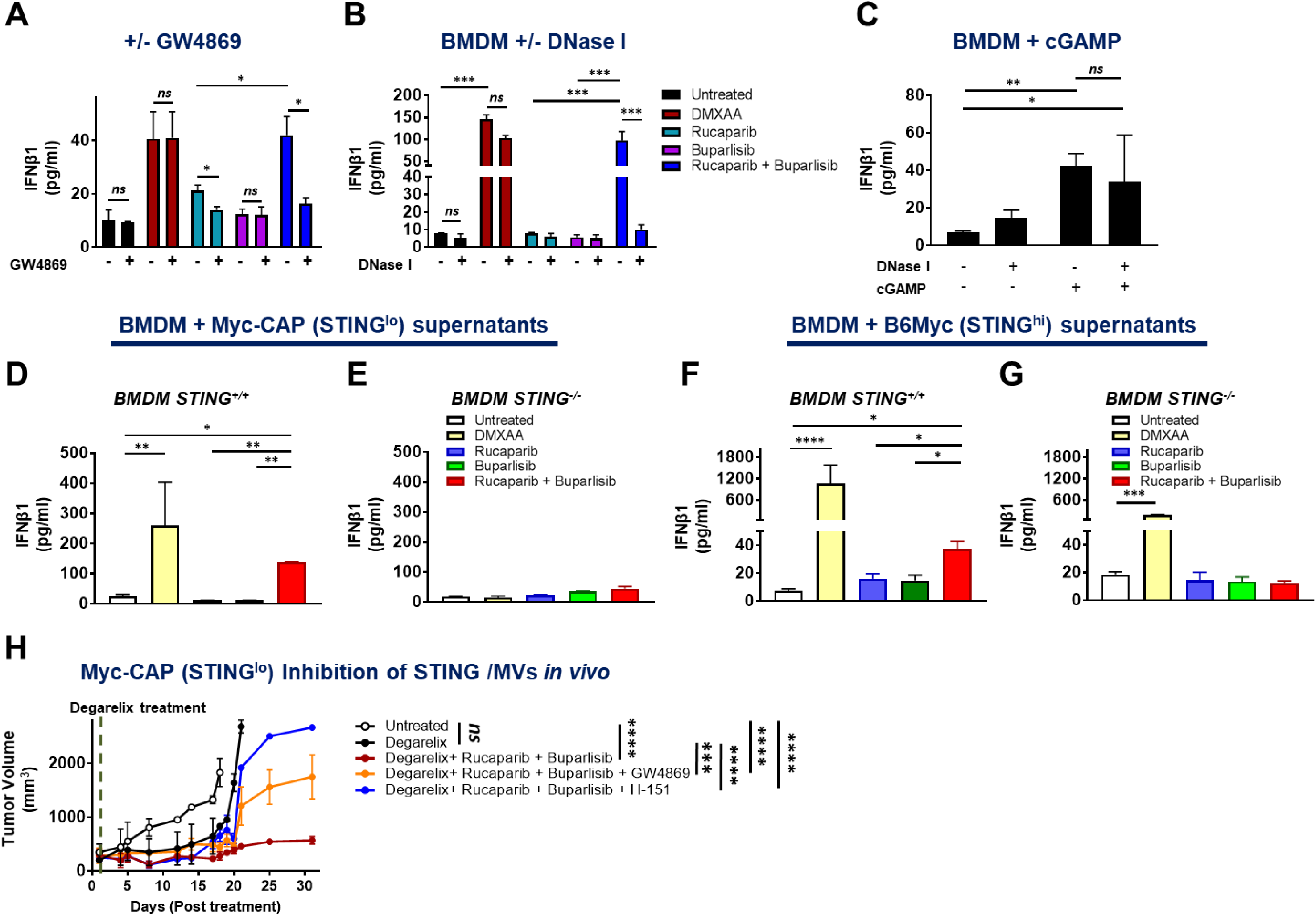
PARPi/PI3K treatment activates tumor-cell extrinsic cGAS/STING pathway within TAMs *in vivo*, via MV surface-associated DNA DSBs released from tumor cells. **(A)** Syngeneic mice were engrafted with Myc-CAP (STING^lo^) cells, and tumors were harvested when the tumor volume reached 400 mm^3^. Single cell suspensions of the tumors were treated with the indicated drug(s), in the presence or absence of GW4869 (MV biogenesis inhibitor). **(B)** Bone marrow derived macrophage (BMDMs) were co-cultured for 30 hours with supernatants (-/+ exogenous 50 units of DNase I pre-treatment for 30 minutes) from Myc-CAP cells that were treated with the indicated drug(s) for 36 hours. **(C)** cGAMP (10 μg/ml) was pre-incubated with 10 units of DNAse I or mock control for 30 minutes and then treated with BMDMs for 30 hours. **(D-G)** STING^+/+^/STING^-/-^ BMDMs were co-cultured for 36 hours with supernatants from Myc-CAP and B6-Myc cancer cells, that were treated with the indicated drug(s) for 36 hours. Supernatants were collected from **A-G** at the end of treatment and analyzed for IFNβ1 by ELISA. **(H)** Myc-CAP (STING^lo^) tumor-bearing mice were treated with the indicated drug(s) until tumors first reached approx. 2500 mm^3^. For the degerelix/rucaparib/buparlisib combination, additional cohorts of mice underwent concomitant treatment with STING antagonist H-151 or GW4869. Tumor volume was recorded daily for duration of experiment. For *in vitro* experiments, n=2 independent experiments and for *in vivo* experiments n= 3-4 animals /group. Significance/p-values were calculated by one-way ANOVA and are indicated as follows, *p < 0.05; **p < 0.01, ***p < 0.001; **** p< 0.0001; ns = not statistically significant;

If DNA DSB fragments are localized to the MV surface resulting in activation of cGAS/STING pathway activation within TAMs, then co-culture of bone marrow derived macrophages (BMDMs) with DNase I treated supernatants from rucaparib/buparlisib-treated cancer cells would be predicted to result in abrogation of Type I IFN response relative to untreated MVs. On the other hand, if DNA DSB fragments are enclosed within MVs, then DNase I treatment would have no effect on induction of Type I IFN response within BMDMs. We observed a striking decrease of IFNβ release from BMDMs that were co-cultured with rucaparib/buparlisib-treated Myc-CAP supernatants that were treated with DNase I, relative to the corresponding supernatants that were not treated with DNase I **(Fig 5B)**, thus demonstrating that MV surface-associated (and not internalized) DNA DSB fragments are responsible for the Type I IFN response elicited within BMDMs.

Recent studies have demonstrated that direct transfer of cGAMP through tight junctions can activate the cGAS/STING pathway within the TME (34). To test the hypothesis that DNA DSBs, and not cGAMP, is responsible for cGAS/STING pathway activation within TAMs, we co-cultured BMDMs with recombinant cGAMP that was pre-treated with DNase I. Importantly, DNase I pre-treatment did not abrogate the Type I IFN production within BMDMs in response to cGAMP **(Fig. 5C)**, demonstrating that it is the MV surface-associated DNA DSBs, not direct cGAMP transfer within the TME, that is responsible for cGAS/STING activation within TAMs.

To confirm that cGAS/STING pathway activation within TAMs is responsible for the Type I IFN response within the TME, supernatants from B6-Myc (STING^hi^) and Myc-CAP (STING^lo^) cancer cells treated with rucaparib and buparlisib, singly or in combination, were co-cultured with STING-proficient and STING-deficient BMDMs **(Fig. 5D-G)**. We observed that the rucaparib/buparlisib-induced IFNβ production observed when conditioned supernatants from B6-Myc (STING^hi^) and Myc-CAP (STING^lo^) cells were co-cultured with STING proficient BMDMs **(Fig. 5D, 5F)**, was abrogated under similar conditions with STING^-/-^ BMDMs **(Fig. 5E, 5G)**. To determine whether the DNA DSB-associated MV release from tumor cells and activation of STING within TAMs is relevant *in vivo*, Myc-CAP (STING^lo^) tumor-bearing mice were treated with the degerelix/rucaparib/buparlisib combination in the presence or absence of STING antagonist H-151 or MV biogenesis and release inhibitor GW4869. Critically, we observed an abrogation of anti-tumor response elicited by degarelix/rucaparib/buparlisib combination in Myc-CAP (STING^lo^) tumor-bearing mice that were concomitantly treated with H-151 or GW4869 **(Fig. 5H)**. Collectively, these *ex vivo* and *in vivo* studies revealed that PARPi/PI3K treatment activates tumor-cell extrinsic cGAS/STING pathway within TAMs via MV surface-associated dsDNA cargo released from Myc-CAP (STING^lo^) tumor cells.

### Optimal STING pathway activation within TAMs requires both MV surface-associated DNA DSBs and PI3Ki-mediated de-repression of cGAS activity

Recent work has shown that AKT can phosphorylate cGAS at Ser-291 and Ser-305, leading to post-translational suppression of its enzymatic activity (35). Furthermore, we observed that concomitant PI3Ki did not induce additive DNA damage with PARPi in Myc-CAP (STING^lo^) cells, suggesting that the addition of PI3Ki elicits a non-tumor cell autonomous DNA-damage independent mechanism for cGAS/STING pathway within TAMs. We hypothesized that the requirement for concomitant PI3Ki treatment to activate DNA-damage induced STING pathway activation within TAMs, stems from its ability to de-repress cGAS enzymatic activity, resulting in increased production of STING ligand 2’3’-cGAMP (cyclic guanosine monophosphate–adenosine monophosphate). To specifically test this hypothesis, we treated Myc-CAP (STING^lo^) cells with rucaparib *in vitro* for 36 hours, followed by isolation of MVs from supernatant (R-MVs). Next, we co-cultured BMDMs with R-MVs in the presence or absence of buparlisib (at target inhibitory concentration of 1 µM; **Supplementary Fig. 15**) for 36 hours and measured levels of intra-cellular cGAMP within macrophages by ELISA. Interestingly, we observed a significant induction of cGAMP only with the combination of buparlisib and MVs derived from rucaparib-treated Myc-CAP (STING^lo^) cells, not corresponding single-agent controls **(Fig. 6B)**. Critically, the combination of buparlisib and R-MVs resulted in downstream activation of STING signaling pathway components **(Fig. 6C)** and a significant increase in IFN-α **(Fig. 6D)** and IFN-β **(Fig. 6E)**, similar to what was achieved with direct treatment of BMDMs with STING agonist DMXAA **(Fig. 6D-E)**. Taken together, these data demonstrate that optimal STING activation within macrophages requires both PARPi-mediated MV surface-associated DNA DSBs and PI3Ki-induced de-repression of cGAS enzymatic activity **(Fig. 6F)**.

**Figure 6:**
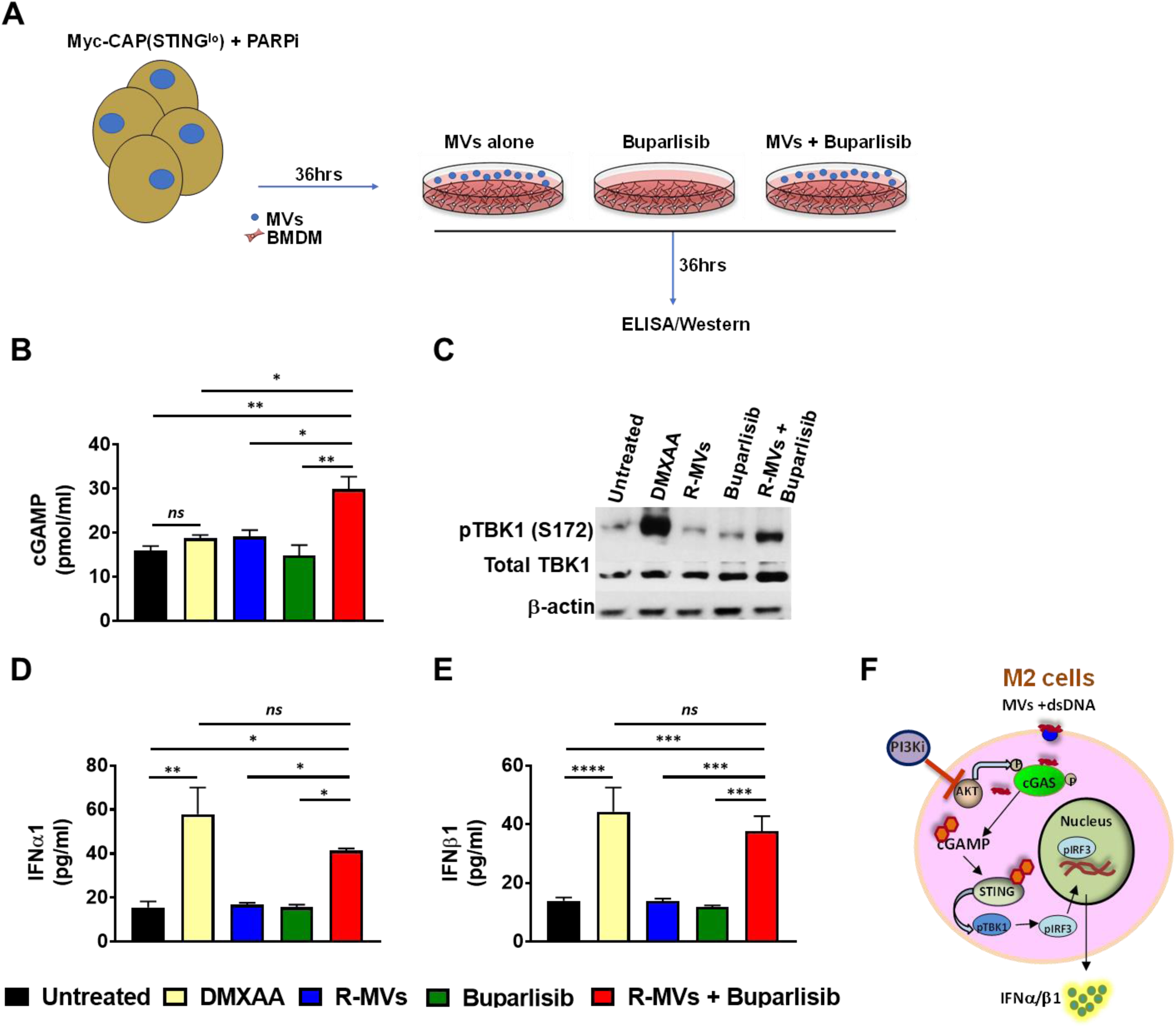
Optimal STING activation within macrophages requires both PARPi-mediated MV surface-associated DNA DSBs and PI3Ki-induced de-repression of cGAS enzymatic activity. **(A)** Experimental schema: Myc-CAP (STING^lo^) cells were treated with rucaparib (0.5 µM) for 36 hours, followed by isolation of MVs from supernatant (R-MVs), which were then co-cultured with BMDMs in the presence/absence of buparlisib (1 µM) for 36 hours. BMDMs were directly treated with DMXAA (50 µg/ml). **(B-C)** Cellular metabolites and proteins extracted from co-cultured BMDMs (as per schema in **A**) were used for cGAMP ELISA **(B)** and for assessment of activation of STING pathway by western blotting using indicated antibodies **(C**), respectively. **(D-E)** Supernatants collected from experiment in **(A)** were used for IFN-α **(D)** and IFN-β ELISA **(E)**. (**F)** Model for cGAS/STING pathway activation within suppressive macrophages (M2) following treatment with buparlisib and MVs (isolated from rucaparib-treated Myc-CAP cells); n=2 independent experiments. Significance/ p-values were calculated by one-way ANOVA and are indicated as follows, *p < 0.05; **p < 0.01, ***p < 0.001, **** p< 0.0001, ns = not statistically significant.

To determine whether PARPi/PI3Ki-induced cGAS/STING pathway activation within macrophages results in their polarization from pro-tumorigenic M2 to anti-tumorigenic M1 phenotype, we co-cultured BMDMs with buparlisib, singly or in combination with purified MVs derived from supernatants of Myc-CAP (STING^lo^) tumor cells that were treated with rucaparib (R-MVs). Flow cytometry analysis revealed an increase in MHC Class II and CD86 expression within macrophages, indicating enhanced activation and antigen presenting capacity, respectively, following treatment with both buparlisib and R-MVs, relative to single agent buparlisib or R-MVs controls. Treatment of BMDMs with DMXAA achieved similar levels of macrophage activation and antigen presentation, relative to buparlisib/R-MVs combination **(Fig. 7A-B)**. Furthermore, we observed a significant increase in TNF-α, and T cell chemoattractant chemokines CXCL10 and CCL5 release **(Fig. 7C-E),** and reversal of CSF-1R and PD-L1 inhibitory marker expression within BMDMs co-cultured with both buparlisib and R-MVs, relative to single-agent controls **(Supplementary Fig. 16A, 16B)**, thus demonstrating M2-to-M1 polarization following R-MVs and buparlisib combination treatment.

**Figure 7:**
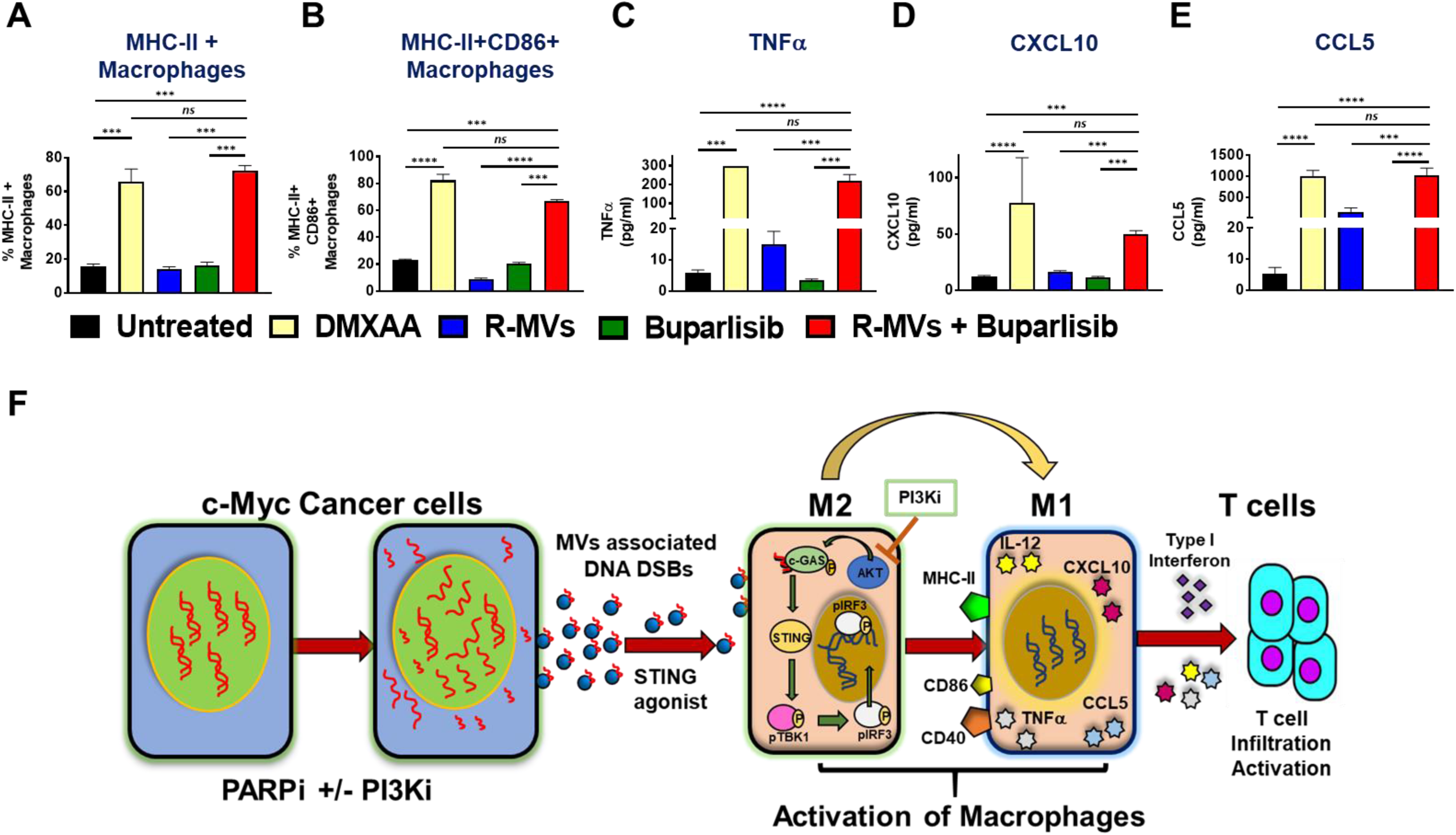
The combination of buparlisib with MV surface-associated DNA DSBs from PARPi treated cancer cells reprograms macrophages from an M2 to M1 phenotype. Myc-CAP (STING^lo^) cells were treated with rucaparib (0.5 µM) for 36 hours, followed by isolation of MVs from supernatant (R-MVs), which were then co-incubated with BMDMs in the presence/absence of buparlisib (1 µM) for 36 hours. As a positive control, BMDMs were directly treated with DMXAA (50 µg/ml). At the end of treatment, BMDMs were analyzed by flow cytometry for macrophage activation markers to quantify frequency of CD45+CD11b+F4/80+ cells expressing MHC-II **(A)** and CD86 **(B)**. **(C-E)** Supernatants were collected for determination of M1-specific cytokines/chemokines by cytokine array; n=2 independent experiments; Significance/ p-values were calculated by one-way ANOVA and are indicated as follow, *p < 0.05; **p < 0.01, ***p < 0.001, **** p< 0.0001, ns = not statistically significant. **(F)** Working Model: The combination of PARPi + PI3Ki induces intracellular DNA DSBs, that become associated with the surface of extracellular MVs following release into the TME. In addition, PI3Ki can inhibit AKT-mediated Ser-291 phosphorylation of cGAS, thus de-repressing its enzymatic activity. MV surface-associated DNA DSBs can secondarily activate the cGAS/STING pathway within TAMs only in the presence of concomitant PI3Ki, resulting in macrophage activation and M2 to M1 polarization, increased T cell infiltration, and tumor regression in c-myc driven PC.

We also performed analogous co-culture experiments of BMDMs with supernatants derived from rucaparib-treated Myc-CAP (STING^lo^) cells, in the presence or absence of buparlisib. Strikingly, flow cytometry analysis revealed increased MHC Class II and iNOS expression, and decreased Arginase I expression only with the rucaparib/buparlisib combination, not corresponding single-agent controls **(Supplementary Fig. 17A-C).** Furthermore, cytokine profiling of supernatants from *ex vivo* experiments revealed an increase in CXCL10 and CCL5 released from BMDMs that were treated with supernatants derived from rucaparib/buparlisib treated Myc-CAP (STING^lo^) cells, not corresponding single agent controls **(Fig. 17D-E)**. Taken together, these results demonstrate that PARPi-induced DNA DSBs and PI3Ki-induced cGAS de-repression within TAMs is required for optimal cGAS/STING activation and re-programming of macrophages from M2 suppressive to M1 activated, anti-tumor phenotype within the TME. A mechanistic model summarizing the findings in this paper is depicted in **Fig. 7F**.

## DISCUSSION

Androgen receptor (AR)-directed therapies have had incremental benefit, but are generally not curative for the treatment of metastatic PC. There has been renewed interest in PC immunotherapy, partly based on the profound and durable clinical responses to ICB antibodies targeting CTLA-4 and PD-1/PD-L1 in other cancers (3, 36). While, *de novo* androgen deprivation therapy (ADT) can induce immune cell infiltration within non-inflamed tumor microenvironment of PC only 10-25% of mCRPC patients, respond to ICB (4–6). Here we show a paucity of immune cell infiltrate within the TME in mCRPC patients, which is one mechanism of resistance to ICB. Furthermore, we demonstrate that the majority of immune cells within the TME of mCRPC patients are comprised of TAMs (Fig. 1). On the basis of these findings, we hypothesized that activation of innate immunity within TAMs will enhance immune-responsiveness in PC. Critically, we demonstrate that targeting both fundamental DNA repair and oncogenic signaling pathways could markedly increase responsiveness to ICB via activation of the cGAS/STING pathway within TAMs.

While PARPi have been FDA approved in BRCA1/2-mutated breast cancer (18), ovarian cancer (19), mCRPC (16, 17) and pancreatic cancer (20), it has limited efficacy in non-BRCA HR-deficient mCRPC (37), as well as HR-proficient-cancers (24). Here we demonstrate that PARPi, either singly and/or in combination with PI3Ki, can induce DNA damage in HR-proficient c-myc driven murine PC models **(Supplementary Fig. 2, 14)**. This is likely related to enhanced dependency of c-myc induced replicative stress on PARP-1 mediated DNA repair (38), with resultant downregulation of the CDK18/ATR axis (39). Consistent with clinical observations, PARPi-induced DNA damage is insufficient to induce apoptosis of HR-proficient c-myc-driven cancer cells *in vitro* **(Supplementary Fig. 2, 14)**.

The cGAS/STING pathway is physiologically activated by cytosolic double-stranded DNA (dsDNA), which typically occurs in the context of viral infections. Cyclic GMP-AMP synthase (cGAS) is a primary cytosolic dsDNA sensor that generates cyclic dinucleotides (cGAMP), which acts as a second messenger to activate STING, which in turn induces the recruitment of TBK1 and IRF-3 to form a complex with STING (40). The activation of IRF-3 and/or NF-κB signaling pathways induce the expression of Type I IFNs and pro-inflammatory cytokines (41, 42). Recent murine studies have demonstrated that PARPi can activate tumor cell-intrinsic cGAS/STING pathway in murine HR-deficient breast and ovarian cancers, resulting in anti-tumor responses that can be accentuated with PD-1 blockade (21, 22). This has led to several clinical trials evaluating radiotherapy or PARP inhibitors with ICB in different solid tumor malignancies.

Our clinical trial data and preclinical studies described here demonstrate that PARPi is insufficient to drive tumor cell-extrinsic cGAS/STING pathway activation and does not enhance ICB responsiveness in a co-clinical trial testing PARPi/ICB combination in HR-proficient mCRPC patients and c-myc-driven murine models of prostate cancer **(Fig. 3-4)**. Critically, concomitant PARPi/PI3Ki treatment activates tumor cell-extrinsic cGAS/STING pathway activation within M2 TAMs, resulting in their polarization into an anti-cancer M1 phenotype, and T cell infiltration/activation and tumor regression in immune-refractory HR-proficient c-myc-driven models of PC **(Fig. 7F)**. This PARPi/PI3Ki-mediated tumor regression *in vivo* is abrogated with systemic macrophage depletion, demonstrating that the reprogramming of TAMs is responsible for driving anti-cancer innate immunity. Furthermore, the PARPi/PI3Ki combination fails to induce apoptosis of B6-Myc (STING^hi^) and Myc-CAP (STING^lo^) cancer cells *in vitro*, and effectively control tumor growth *in vivo* in corresponding immunodeficient models. Collectively, these findings demonstrate that PARPi/PI3Ki combination exerts its anti-cancer activity primarily via a non-tumor cell autonomous, innate immune, macrophage-driven mechanism. Given this unanticipated immune-based mechanism for PARPi/PI3Ki combination in c-myc driven PC, these data highlight the critical unmet need for the development of more sophisticated *ex vivo* and *in vivo* combinatorial drug screening platforms, which incorporate immunological readouts beyond apoptosis induction of cancer cell lines *in vitro*.

Consistent with prior studies that have shown that PARPi, in combination with PI3Ki, induces additive DNA damage and suppresses tumor growth in breast and prostate preclinical models (28, 43), we observed additive DNA damage in B6-Myc (STING^hi^) tumors with rucaparib in combination with buparlisib. In contrast, the addition of buparlisib to Myc-CAP (STING^lo^) cells did not further increase DNA damage, relative to rucaparib alone, thus suggesting that concomitant PI3Ki treatment elicits anti-cancer effects via a DNA-damage independent mechanism in Myc-CAP (STING^lo^) tumors. A recent study has demonstrated that AKT phosphorylates the S^291^ or S^305^ of the carboxyl-terminal enzymatic domain of mouse or human cGAS, respectively, and that this phosphorylation robustly suppresses cGAS enzymatic activity, leading to decreased cytokine production and antiviral activity following DNA virus infection (35). Furthermore, our *ex vivo* studies revealed that presence of DNA DSBs within the TME is insufficient to drive cGAS/STING pathway activation within TAMs. We therefore tested the hypothesis that the PI3K/AKT pathway suppresses c-GAS enzymatic within TAMs, thereby preventing STING pathway activation in response to PARPi-induced DNA DSBs. Consistent with this hypothesis, we observed that PI3Ki de-represses cGAS enzymatic activity, resulting in increased cGAMP production, STING activation and Type I IFN production, which is required for DNA DSB induced cGAS/STING activation within TAMs.

In this study, we have made the exciting observation that PARPi-induced DNA DSBs are transported within the TME as cargo associated with the surface of MVs, which can secondarily activate cGAS/STING pathway in TAMs. Furthermore, this DNA DSB-induced cGAS activation within TAMs is abolished by concomitant DNase I treatment, suggesting that the DNA DSB is associated on the surface of the MVs, and not internally within the membrane lipid bilayer. These findings are supported by a recent study has shown that dsDNA can be associated with the surface of exosomes (44). Given that exosomes/MVs can have immunosuppressive and pro-metastatic properties (45, 46), our findings suggest the possibility that PARPi can render MVs more immunogenic, similar to prior observations made with MVs produced after radiotherapy (47). Future studies will be needed to evaluate the possibility of utilizing blood-based MVs biomarkers as pharmacodynamic readouts of PARPi-induced DNA damage within the TME.

Prior studies have demonstrated that early stages of castration induce T-effector and T-regulatory cell infiltration within human and prostate tumors (32, 33). In this study, we interrogated two different murine models of c-myc-driven PC, B6-Myc (STING^hi^) and Myc-CAP (STING^lo^), which express high and low levels of cGAS/STING signaling pathway components **(Supplementary Fig. 1)**, respectively, and mimic the heterogeneity of cGAS and STING expression observed in human PC **(Fig. 2)**. In the B6-Myc (STING^hi^) context, the combination of PARPi/PI3Ki was sufficient to drive a macrophage-mediated innate immune response and tumor clearance, whereas Myc-CAP (STING^lo^) bearing syngeneic mice were *de novo* resistant to this combination. Strikingly, we observed that the addition of ADT, when combined with PARPi/PI3Ki, resulted in an anti-cancer innate immune response and tumor clearance, similar to that observed in B6-Myc mice treated with the combination without castration. There are several potential explanations for these findings. First, C57BL6 is a more immunogenic strain than FVB mice (29, 31), resulting in a lower threshold for immune-sensitization that is dependent on host factors. Second, the presence of higher tumor cell intrinsic STING levels in B6-Myc, relative to Myc-CAP cells, could account for the approx. 10-fold higher baseline CD45+ immune cell infiltration in B6-Myc tumors in vivo, relative to Myc-CAP tumors. Following castration, Myc-CAP (STING^lo^) tumors have an approx. 5-fold increase in CD45+ immune cell infiltration (predominantly TAMs), which provides the necessary immunological milieu needed for optimal cGAS/STING activation within the TME following PARPi/PI3Ki treatment. Collectively, these data suggest that baseline cGAS and STING expression can be developed as potential biomarkers for response to PARPi/PI3K combination therapy in earlier stages of PC treatment, where castration is not standard-of-care. Given our findings that high-risk (Gleason ≥8) PC is enriched for tumors with low tumor cell intrinsic STING expression, the findings in this paper warrant the development of immuno-oncology clinical trials testing the combination of ADT with PARPi/PI3Ki/PD-1 blockade in *de novo* hormone-sensitive, locally advanced or metastatic PC.

In summary, we have demonstrated that concomitant targeting of PARP and PI3K signaling pathways can trigger non-tumor cell autonomous c-GAS/STING pathway activation within TAMs, thereby enhancing T cell recruitment/activation into the TME and tumor regression in HR-proficient c-myc-driven murine models. Based on these findings, PARPi/PI3Ki combination therapy could markedly increase the fraction of PC patients responsive to ICB, independent of HR status, and clinical trials to test this combinatorial approach are warranted.

## METHODS

### Rucaparib/nivolumab clinical trial in mCRPC patients

mCRPC prostate cancer patients, independent of HR status, who had received at least one AR targeted therapy, without prior exposure to PARPi or ICB therapy, were enrolled in an investigator-initiated, IRB-approved co-clinical trial (NCT03572478) of rucaparib (PARPi) with nivolumab (PD-1 antibody), that was co-sponsored by Clovis Oncology and BMS respectively. The patients were treated until disease progression or unacceptable toxicity. All patients provided informed consent prior to clinical trial enrollment. As part of study requirements, serial PSAs were obtained on a monthly basis following study enrollment and measured using standard clinical laboratory diagnostic methods.

### Multi-Parameter Flow Cytometry

#### Human Biopsies

Tissues were processed into single cell suspensions via gentle mechanical dissociation in 12-well plates containing 1 ml of 10% RPMI media supplemented with 10% fetal bovine serum, 1% penicillin-streptomycin and 2% L-glutamine. Cell suspensions were centrifuged at 500g for 5 minutes at 4 °C, resuspended in FACS buffer (1X PBS containing 0.5% FBS and 0.01% sodium azide) and used for staining with the following anti-human antibodies (Biolegend): CD45, CD11b, CD163, CD68, HLA-DR, CD15, CD33, CD16, PD-L1, CD3, CD4, CD8 CD19, 4-1BB, PD-1, CD11c. All flow antibodies in this study were utilized at recommended dilutions provided by the manufacturer.

#### Murine tumors

Murine tumors were processed identically with an additional step of filtration to remove cell debris, where single cells were passed through a 70-micron mesh, prior to stain. One million cells resuspended in 1X FACS buffer were stained with titrated concentrations of the following anti-mouse antibodies: CD45, CD11b, CD11c, CD19, F480, Ly6G, Ly6C, PD-L1, and VISTA, I-A^e^/IA^b^, H-2^Kb^, CD3, CD4, CD8, 4-1BB, PD-1, CD206, CSF-1R. Incubation with antibodies was done at 4 °C for 30-40 minutes for both murine and human cells. Following staining, cells were washed twice with IX FACS buffer and fixed with 300 μl of 4% paraformaldehyde (Fisher Scientific), prior to analysis on BD instrument LSR 4-15 Fortessa. Data collected on flow cytometer using BDFACSDIVA software and was analyzed using Flow Jo software (Tree Star).

### Bioinformatics analysis of myeloid gene signature using TCGA database

The transcriptome data (Illumina HiSeq RNASeqV2) was downloaded for prostate tumors and normal prostate from the TCGA data portal (http://tcga-data.nci.nih.gov/tcga/tcgaHome2.jsp) and analyzed for differential expression of STING and c-GAS. Additionally, the Biochemical Recurrence (BCR) status and Gleason Scores were also downloaded for 488 prostate tumors. The samples were grouped into High Gleason score (≥ 8) and Low Gleason score (6/7). The RNA-Seq data was used to analyze the differential expression of genes between high and low Gleason score samples. Statistical analysis evaluating changes in gene expression between the different groups were done using unpaired non-parametric t-test.

### Cancer Cell Lines

Transgenic c-myc^hi^ prostate tumor derived cell line, Myc-CAP (STING^lo^) (25) was obtained from ATCC and passaged in 1X DMEM (without phenol red) containing 10% fetal bovine serum, 1% penicillin-streptomycin and 2% L-glutamine. The corresponding c-myc^hi^ line derived from a C57BL6 generated background (26) were grown for *in vitro* studies, using same culture conditions as for Myc-CAP. All cell lines were confirmed to be Mycoplasma-free, using Universal Mycoplasma Detection Kit (ATCC® 30-1012K™) testing kit. Both B6-Myc (STING^hi^) cell line and B6-Myc whole tumor explants used for *in vivo* studies were a kind gift from Dr. Leigh Ellis (Dana Farber Cancer Institute, Boston). For *in vitro* drug treatments, the following concentrations were used: Rucaparib (500 nM), Buparlisib (1 μM), DMXAA (50 μg/ml) with specific treatment durations for individual experiments indicated in the figure legends.

### Western Blot Analysis

RIPA and T-PER buffer (Thermo Scientific), supplemented with protease (Roche) and phosphatase inhibitor cocktail (Roche), were used for preparation of lysates from *in vitro* cell lines and whole tumor chunks, respectively. For western blotting, the following antibodies were used from Cell Signaling Technology: Polyclonal rabbit anti-mouse-phospho-γH2AX, phospho-AKT, total AKT, cGAS, STING, phospho-IRF3, total IRF3, phospho-TBK1, total TBK1, PTEN, β-actin and GAPDH. Monoclonal anti-mouse PAR antibody was obtained from Trevigen. Images of scanned blots were processed using ADOBE Photoshop.

### Generation of BMDMs

Bone marrow derived macrophages were differentiated as previously described (48). Briefly, bone marrow cells were isolated from male FVB/NJ, C57Bl/6J^STING+/+^ and C57BL/6J-Sting1^gt^/J^(STING-/-)^ mice and differentiated in the presence of 10X RPMI media (supplemented with 10% fetal bovine serum, 1% penicillin-streptomycin and 2% L-glutamine) containing 30% L-conditioned media or M-CSF (50 ng/ml) for 5-7 days. stimulated directly with 50 μg/ml of 5,6-Dimethylxanthenone-4-acetic Acid (DMXAA, mouse STING agonist) for 36 hours. Following treatment, supernatants were collected for Type I IFN ELISA (LEGEND MAX^TM^ Mouse IFNβ1 ELISA, Biolegend) and processed as specified in protocol.

### Generation of syngeneic models and in vivo drug administration

Wild-type (WT) C57BL/6J, C57BL/6J-Sting1^gt^/J^(STING-/-)^, FVB/NJ mice, Athymic nude (Nu/J) and NOD-SCID(NOD.CB17-Prkdc /J) were purchased from Jackson laboratories and mice were kept in an AALAC (American Association for the Accreditation of Laboratory Animal Care) certified barrier facility at the University of Chicago. Animal work was carried out according to approved Institutional Animal Care and Use Committee protocols. For Myc-CAP-based experiments, mice aged 8-10 week were engrafted with 1 million Myc-CAP cells re-suspended in 1X PBS, under anesthesia. For experiments using B6-Myc, 5 mm^2^ tumor chunks were implanted subcutaneously in mice. Treatments were started when tumor volumes reached approximately 200-400 mm^3^, and mice were randomly allocated to treatment groups as indicated. For *in vivo* treatments, lyophilized drugs were reconstituted in appropriate solvents and were administered at the following doses: Degarelix (0.625 mg/kg) was administered as a single intraperitoneal (i.p.) injection. Rucaparib (Clovis Oncology) and buparlisib (the Stand up to Cancer Drug Formulary at Dana Farber Cancer Institute) were administered daily by oral gavage at 150mg/kg and 30 mg/kg, respectively, whereas anti-mouse PD-L1 (clone 10F.9G2; BioXcell) was administered i.p. at 100 µg once every 2 days. DMXAA was injected intratumorally once at a dose of 500 µg/kg. Exosomal Inhibitor GW4869 (Sigma Aldrich) and STING antagonist H-151 (Invivogen) were dosed at 500 μg/gm of body weight i.p. daily and 750 nanomoles/kg i.p. daily, respectively. For *in vivo* macrophage depletion studies, Clodronate (Standard Macrophage Depletion kit, Encapsula Nanosciences) was injected i.p. on a weekly basis at recommended dose of 300 μl of clodronate-liposomal emulsion containing 18.4 mM concentration of clodronate). All *in vivo* treatments were done for 15-28 days and tumor volume measurements were collected on a daily basis. Tumor volume was calculated using the formula: 0.5 × longest diameter × (shortest diameter)^2^. Euthanasia was performed for mice bearing tumor ulceration and/or tumor diameter >2 cm, as per IACUC-approved protocol.

### Confocal Microscopy

Tumor cells were grown at titrated seeding density in glass bottom plates and treated with indicated drug(s) at concentrations described above. Following 36 hours of treatment, culture media was aspirated and the cells were washed twice with 1X PBS. Cells were then fixed with 4% paraformaldehyde at 4 °C, followed by permeabilization briefly with cold 100% ethanol for 8 mins at 4 °C. Staining for DNA DSBs was done with anti-mouse primary antibody, specific for phospho-γH2AX (1:500 dilution, Cell signaling) and secondary anti-rabbit IgG antibody conjugated to AF647 (1:1000-1:2000 dilution in 1X PBS, Thermo Fischer Scientific). Anti-mouse specific β-actin conjugated to Phycoerythrin (PE, Thermo Fischer Scientific) was used to stain the cytoskeleton. All staining procedures were done at 4 °C for 30 minutes. Cells were then washed 3 times with IX PBS and imaged immediately. All images were collected using an Olympus Fluoview 1000 using a 100X oil objective. Acquired images were analyzed by Image J software, developed at NIH.

### MV Isolation/DNA extraction

Cells were treated for 36 hours with the indicated drug(s), and supernatants were harvested and then centrifuged at 300g for 5 minutes at 4 °C to pellet cells. This was followed by additional centrifugation steps at 2,000g for 10 min at 4 °C to eliminate dead cell debris and at 10,000g for 30 min in at 4 °C to remove larger vesicles. The supernatant was then collected and subjected to 100,000 g centrifugation in a Type 60 Ti rotor (38000 rpm) for 70 min at 4 °C. The 100,000g pellet was suspended in 1X PBS to the initial volume of supernatant (2 ml), and washed by an additional spin in the ultracentrifuge for 70 min at 4 °C. The final MV pellet was collected in 1X PBS and used for quantification. Measurements of particle size distribution (PSD) and concentration were performed with a Nanosight LM10 HS-BF instrument (Nanosight Ltd, UK), based on NTA measurements, using a 405-nm 65-mW laser and an EMCCD Andor Luca camera, and revealed MV size range of 50-100 nm.

Samples were diluted with particle-free PBS in a 1:100 dilution (pH = 7.4) to reach the optimal concentration for NTA. All measurements were performed under the identical camera settings (Shutter: 850, Gain: 450, Lower Threshold: 910, Higher Threshold: 11180, 60 s) and processing conditions (NTA 2.3 build 0033, Detection Threshold: 9 Multi, min Track Length: Auto, min Expected Size: minimum of 30 nm). Measurements were performed in multiple repeats (n=3) to collect an at least 5000 events, and then equal numbers of MVs were used for downstream assays. Equal numbers of MVs were collected from each treatment group and DNA was extracted using Trizol LS as per protocol (Thermo Fischer Scientific), and then quantified using Nanodrop.

### Viability Assays

Single cell suspensions at a concentration of 0.5 million cells/0.5ml of 1X Annexin Buffer, were stained with Annexin V/PI (FITC Annexin V Apoptosis Detection kit, BD Biosciences) as specified in manufacturer protocol. Acquisition and analysis of the data sets were done as previously described in section on Multi-parameter flow cytometry.

### Ex vivo reconstitution assay

Subcutaneous tumors were isolated by gentle mechanical dissociation in the presence of 10X RPMI (supplemented with 10% fetal bovine serum, 1% penicillin-streptomycin and 2% L-glutamine). 1X ACK was used for RBC lysis. Single cell suspensions were washed twice with media, by centrifuging at 500g for 5 minutes at 4 °C and quantified using 0.1% Trypsin solution. For sorting of CD45+ cells, PE selection kit (EasySep™ Mouse PE Positive Selection Kit) was used for staining and magnetic extraction of positively labeled cells, as per protocol from vendor. Single cells suspensions derived from tumor were seeded at a concentration of 0.2-0.5 million cells/ml and treated with the following drugs: DMXAA (50ug/ml), rucaparib 500 nM; buparlisib at 1000 nM, singly or in combination, in the presence or absence of exosome inhibitor GW4869 (7.8ng/ml), for 36 hours. All drug stocks were reconstituted in DMSO and further diluted in media used for cell lines *in vitro*. Supernatants were collected at the end of 36 hours and processed as per ELISA protocol for detection of Type I Interferon, as described above.

### Ex Vivo Co-culture Studies with BMDMs

For DNAse I studies, Myc-CAP cells were treated with PARPi or PI3Ki, singly or in combination for 36 hours, and supernatants were treated with 50 units of DNase I in 1X reaction buffer with MgCl_2_, and incubated at 37 °C for 30 min. For neutralization of the DNase I reaction, 1 µL of 50 mM EDTA was added to the mix and then incubated at 65 °C for 10 min. Following this step, supernatants -/+ DNase I were added to BMDMs for 30 hours, and IFNβ1 secretion was assessed, as described above. To rule out cGAMP as the mediator of STING pathway activation within BMDMs, 10 μg of cGAMP disodium salt (MedChem Express) was reconstituted in RNA/DNAse free water and pre-treated with 10 units of DNase I. For BMDM STING validation studies, STING^+/+^/STING^-/-^ BMDMs were co-cultured for 36 hours with supernatants from Myc-CAP and B6-Myc cancer cells, that were treated with the indicated drug(s) for 36 hours. Supernatants were collected at the end of treatment and analyzed for IFNβ1 by ELISA.

For MV reconstitution studies, Myc-CAP (STING^lo^) cells were treated with rucaparib (0.5 µM) for 36 hours, followed by isolation of MVs from supernatant (R-MVs), which were then co-cultured with BMDMs in the presence/absence of buparlisib (1 µM) for 36 hours. Cellular metabolites and proteins extracted from co-cultured BMDMs were used for cGAMP ELISA and for assessment of activation of STING pathway by western blotting. Supernatants were collected at the end of 30-36 hrs and used for detection of Type I IFN/related cytokines and chemokines by cytokine array.

### cGAMP assay

For the colorimetry-based detection of cGAMP production in M2 macrophages, cells were treated with MVs isolated from rucaparib-treated Myc-CAP cancer cells, in the presence or absence of buparlisib. Following 30 hours of treatment, the cells were harvested and cell lysates processed using recommended buffers (cGAMP detection kit, Cayman Chemical) and then used for incubation with anti-cGAMP antibody and detection conjugate for 2 hours or overnight at 4 °C. Next, substrate was added and cGAMP was detected at the indicated wavelength, as per manufacturer’s instructions

### Quantitative Reverse Transcriptase-Polymerase Chain Reaction (qRT-PCR) for Cytokine/Transcription factor

Snap frozen tumor chunks from *in vivo* treatment groups were used for isolation of RNA using Qiagen RNeasy Plus isolation kit (Qiagen), and then used for RT-mediated cDNA synthesis (cDNA RT kit, BioRad), following which PCR was performed with primers specific for *Il-12b* and β-actin, using SyBr Green Universal master mix (BioRad). Each murine sample was analyzed in triplicate on ViiA™ 7 Real-Time PCR System (Applied Biosystems®). Data generated was normalized to β-actin.

### Intracellular Staining for Arginase I

Single cells isolated from murine tumor/differentiated BMDMs were processed using BD Cytofix/Cytoperm solution kit (Fischer Scientific), as per specified protocol. Anti-mouse Arginase I (R&D Systems) was used at recommended dilution for staining of permeabilized cells for 40 mins at room temperature. For flow cytometry, cells were washed twice with FACS buffer by centrifuging at 500g for 5 minutes at 4 °C and resuspended in 300 μl of FACS buffer.

### Statistical Analysis

One-way ANOVA/Mann Whitney/Unpaired t-test/Paired t-test as well as Kolmogrov Smirnov test was used for used for statistical evaluation of experimental datasets. The specific statistical tests used for individual experiments are specifically indicated within the figure legends.

## ACKNOWLEGEMENTS

We thank Dr. Thomas Gajewski for providing helpful suggestions for this manuscript.

## SUPPLEMENTARY FIGURE LEGENDS

**Supplementary Figure 1:**
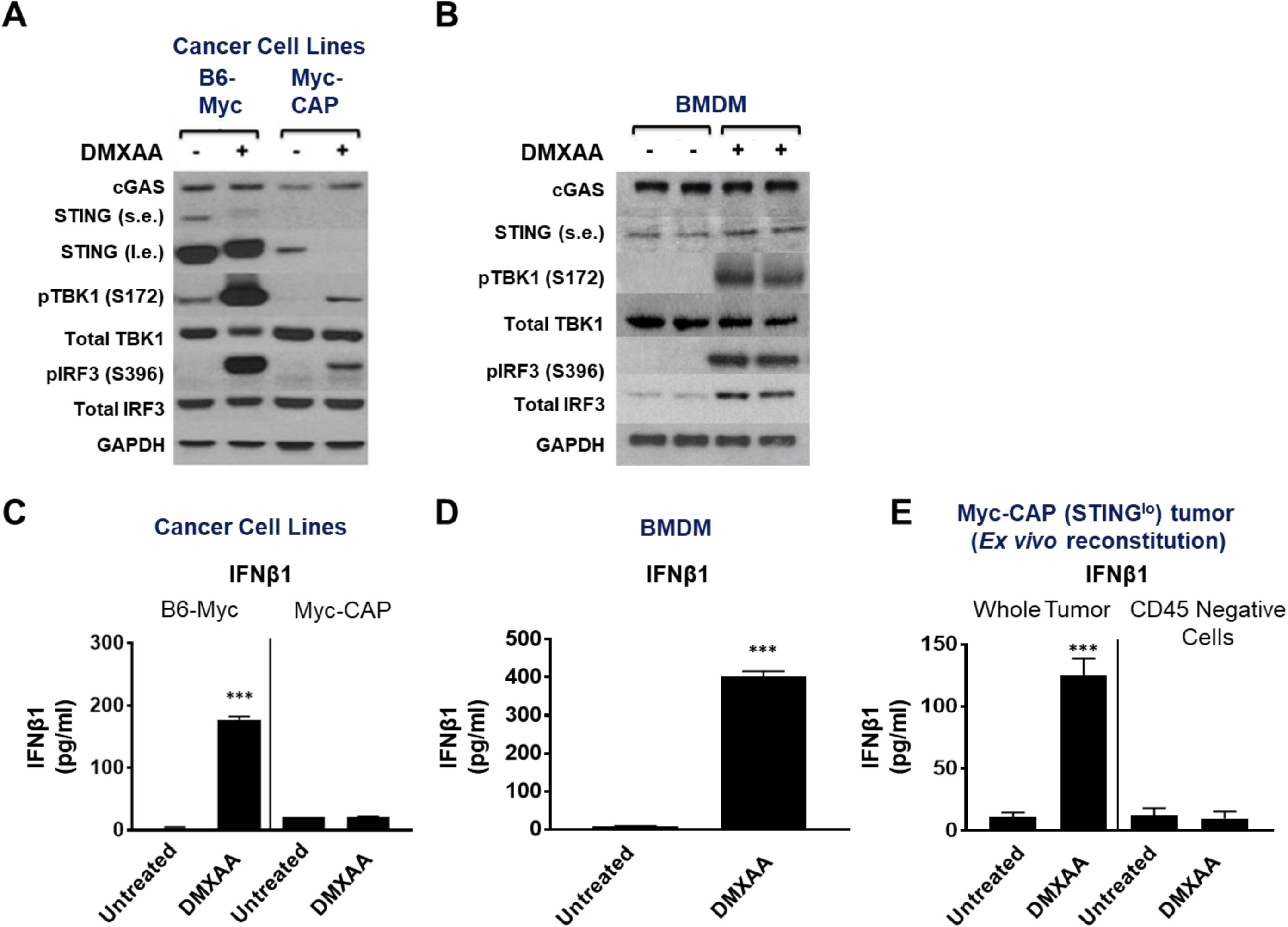
Differential STING pathway activation within B6-Myc (STING^hi^) cell lines and BMDMs, relative to Myc-CAP (STING^lo^) cells. **(A, B)** The indicated cancer cell lines and BMDMs were treated with DMXAA for 1 and 24 hours, respectively and protein extracts were interrogated for cGAS/STING pathway activation with the indicated antibodies by Western blotting. Supernatants were collected for IFNβ1 ELISA, after the indicated cells were treated with DMXAA for 36 hours **(B, D). (E)** Syngeneic Myc-CAP (STING^lo^) tumors were processed into single cell suspensions and used for phycoerythrin-based sorting of CD45+ cellular fractions. Equal number of cells from whole tumor and CD45 Negative fractions were stimulated with DMXAA. Supernatants were collected at 36 hrs for detection of IFNβ1 by ELISA; n=3 independent experiments; Significance/p-values were calculated by one-way ANOVA (C& E), paired t-test (D) and indicated as follows, ***p < 0.001; TAM = tumor associated macrophages; s.e. = short exposure; l.e. = long exposure

**Supplementary Figure 2:**
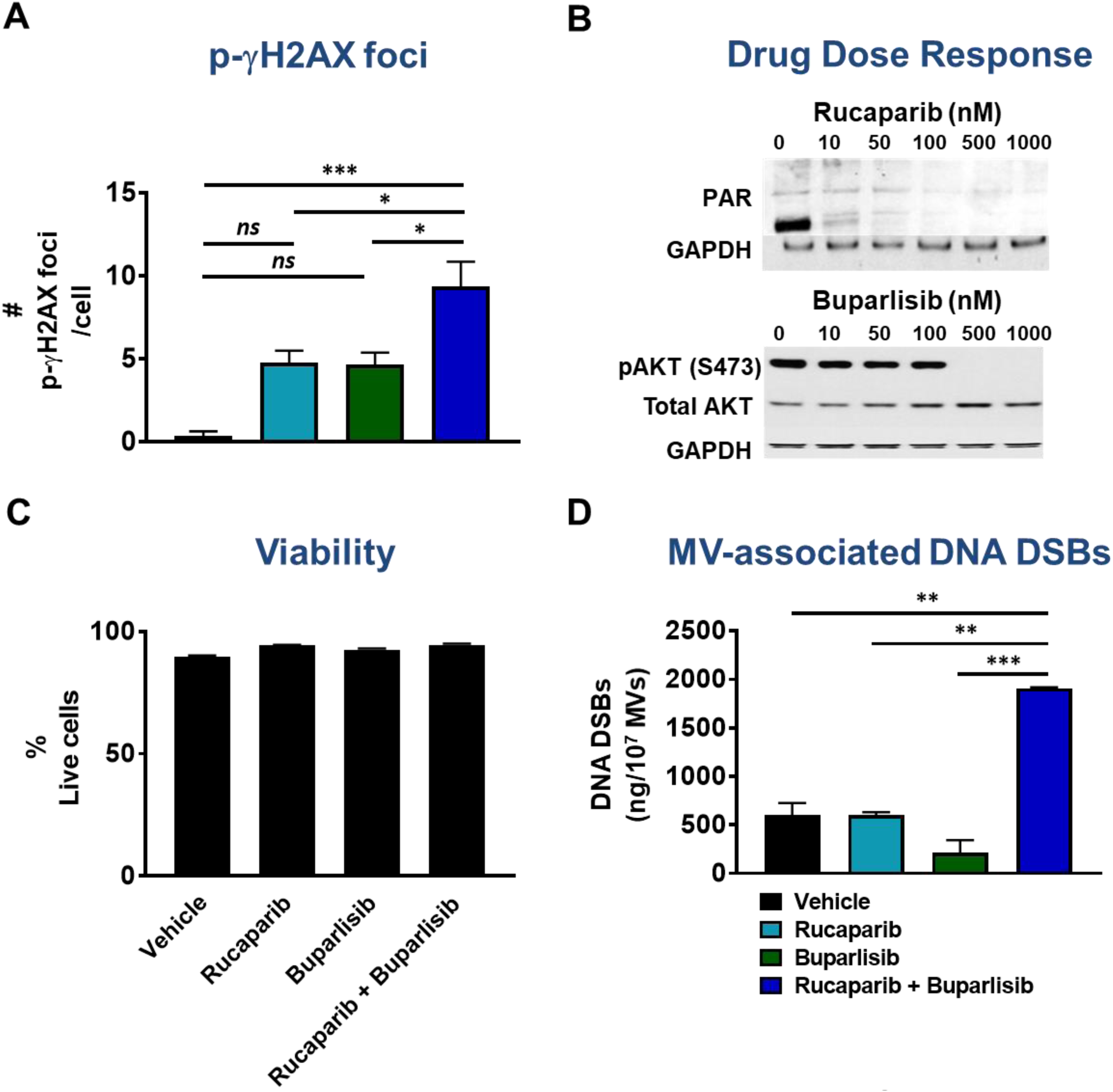
PARPi/PI3Ki combination induces intracellular DNA DSBs, and proportionate release of MV surface-associated DNA DSBs from B6-Myc (STING^hi^) cancer cells, without affecting cellular viability. B6-Myc (STING^hi^) cells were treated with indicated drugs singly or in combination for 36 hours. **(A)** Cells were stained with anti-mouse specific p-γH2AX antibody and fluorescently labeled secondary antibody for determination of DNA DSBs, which were quantified by confocal microscopy. **(B)** Protein extracts from cells in **(A)** were analyzed for the indicated pharmacodynamic biomarkers by western blotting. **(C)** Annexin V-PI staining was done to assess frequency of live cells (Annexin V^-^ PI^-^) following drug treatment. **(D)** Ultracentrifugation was utilized to purify MVs from supernatants in indicated treatment groups **(A),** and associated DNA DSBs was quantified by Nanodrop; n= 2 independent experiments. Significance/p-values were calculated by one-way ANOVA and indicated as follows *p < 0.05; **p < 0.01; ***p < 0.001.

**Supplementary Figure 3:**
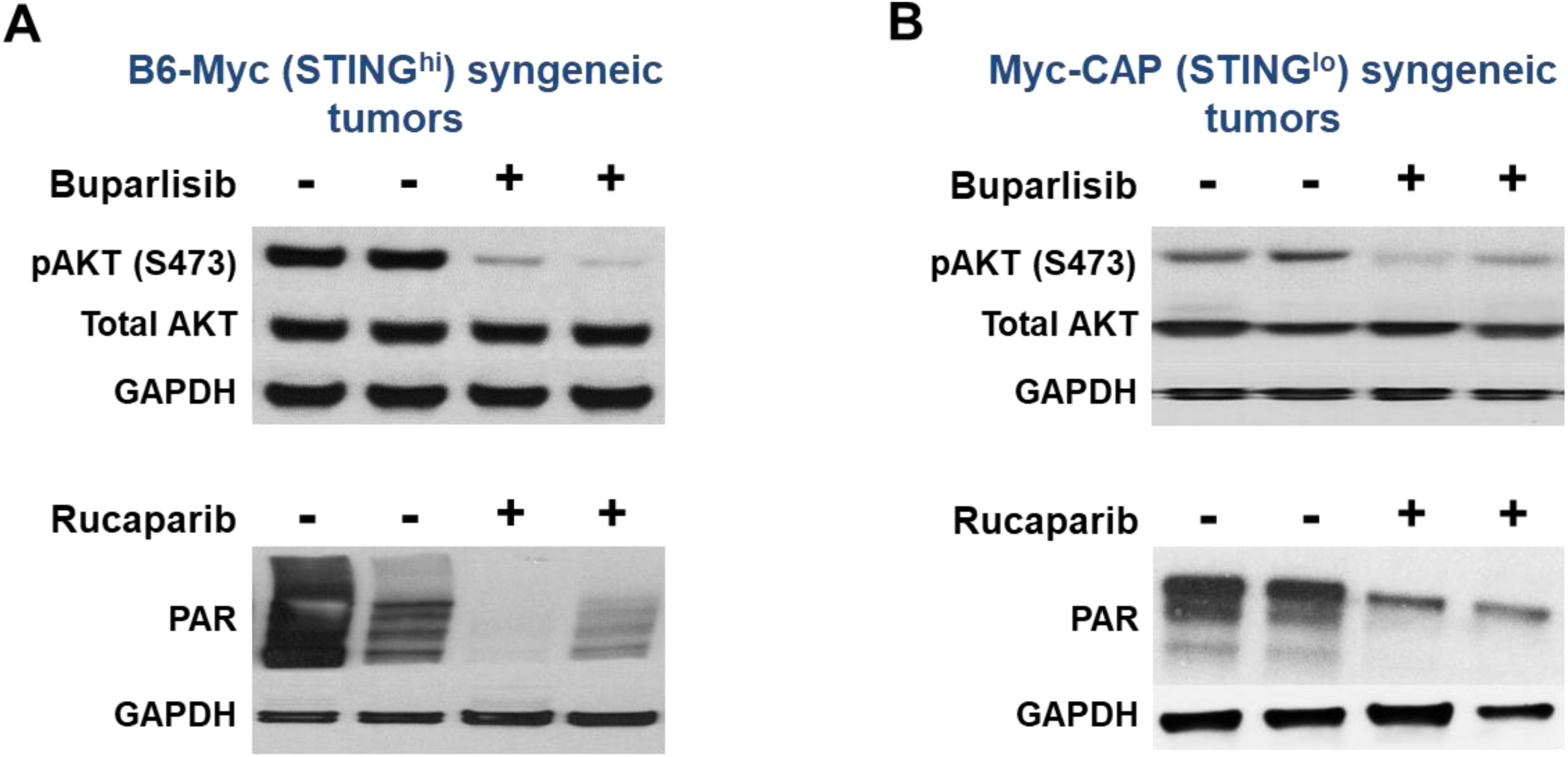
Buparlisib and Rucaparib inhibit intratumoral Pi3K and PARP enzymatic activity, respectively, within B6-Myc (STING^hi^) and Myc-CAP (STING^lo^) tumors. Subcutaneous Myc-CAP (STING^lo^) and B6-Myc (STING^hi^) tumors at 400-500mm^3^ volumes were treated with either buparlisib (pan-PI3K inhibitor, 30mg/kg) or rucaparib (PARP inhibitor, 150mg/kg) by oral gavage daily for 7 days. At end of treatment, both **(A)** B6-Myc (STING^hi^) and **(B)** Myc-CAP (STING^lo^) tumors were harvested and protein extracts utilized for assessment of target inhibition with the indicated antibodies by Western blotting. n=2 mice per treatment group.

**Supplementary Figure 4:**
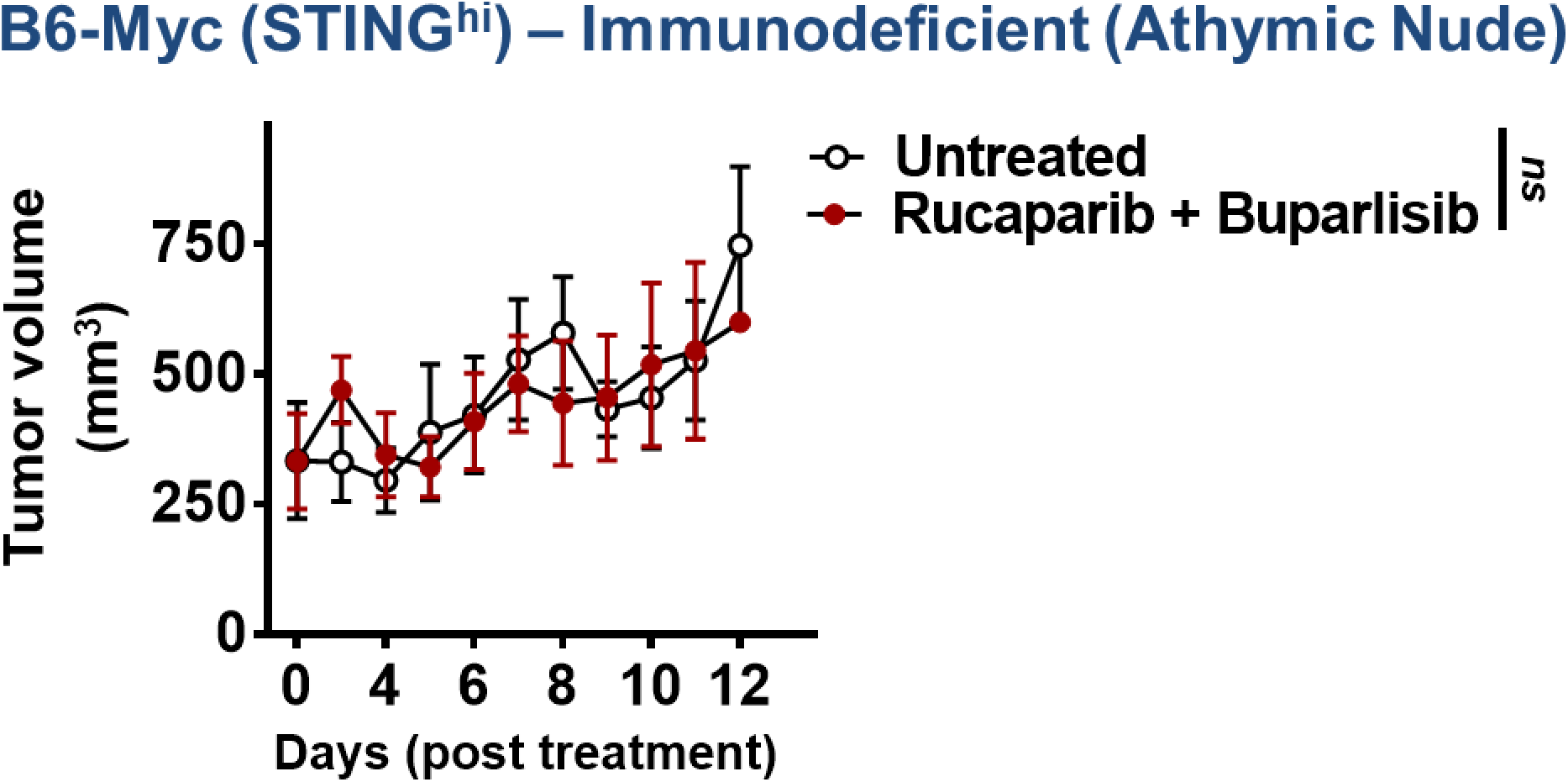
The anti-tumor response elicited by PARPi/PI3K is abrogated in B6-Myc (STING^hi^)-tumor bearing athymic nude mice. B6-Myc (STING^hi^) tumor-bearing athymic nude mice were treated with the indicated drug(s) for approximately 12 days. Tumor volume curves for duration of treatment are shown; n= 3 animals/group; Significance/p-values were calculated by one-way ANOVA and indicated as follows: ns = not statistically significant.

**Supplementary Figure 5:**
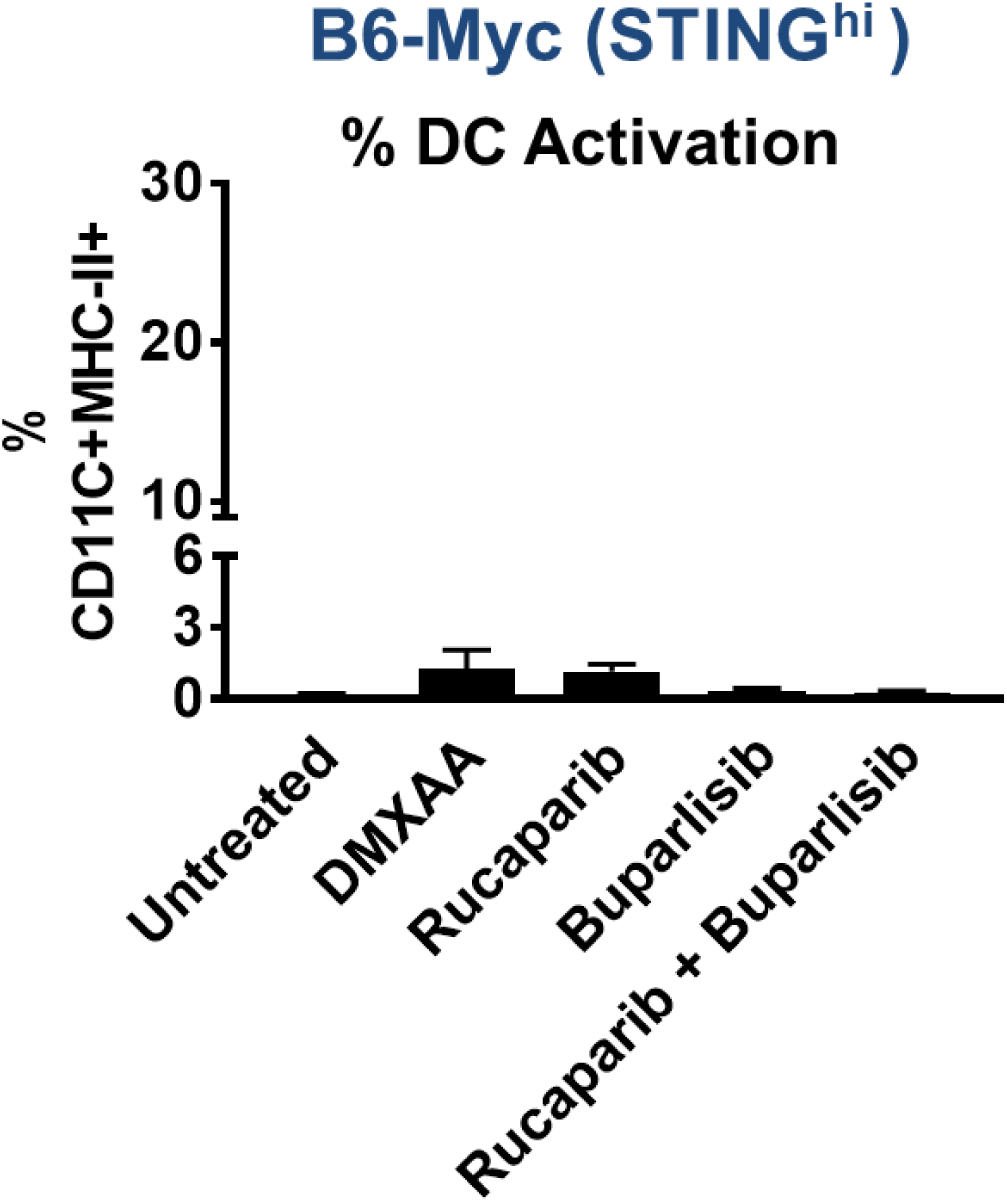
PARPi/PI3Ki-mediated tumor regression in B6-Myc (STING^hi^) model occurs via a DC-independent mechanism. B6-Myc (STING^hi^) tumor-bearing syngeneic mice were treated with indicated drugs(s) until untreated tumors reached approx. 2500 mm^3^. Single-cell suspensions were generated from harvested tumors and analyzed by flow cytometry for % activated DCs gated on CD45+CD11b+ MHC-II+ CD11C+. Data are represented relative to CD45+ immune cells. n=3-4 per group.

**Supplementary Figure 6:**
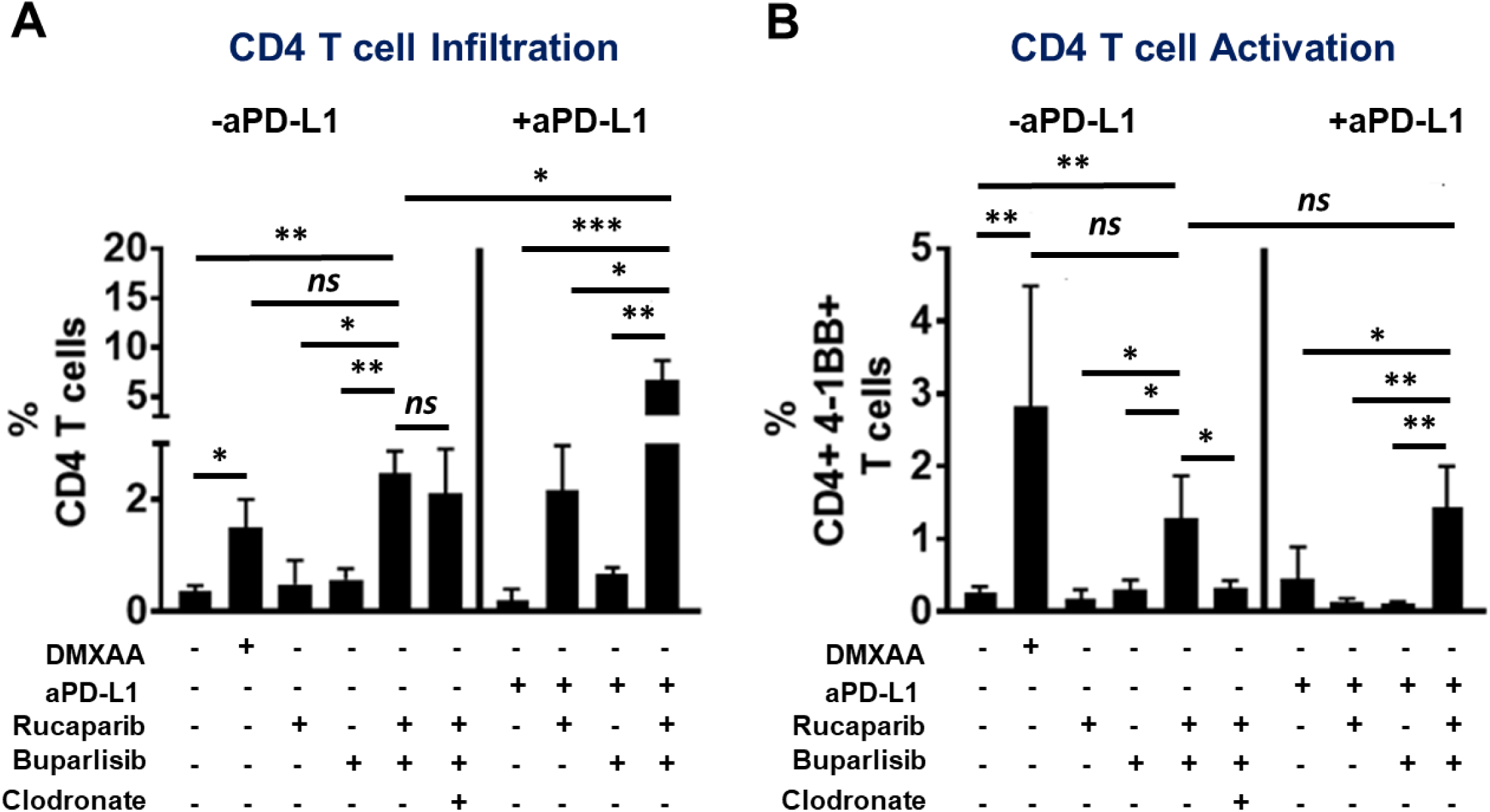
PARPi/PI3Ki/PD-L1 treatment increases CD4 T cell infiltration within the TME of STING^hi^ B6-Myc tumors. B6-Myc (STING^hi^) tumor-bearing syngeneic mice were treated with the indicated drug(s) until untreated tumors reached approx. 2500 mm^3^. Single-cell suspensions were generated from harvested tumors and analyzed by flow cytometry for CD4 infiltration **(A)** and **(B)** activation. Data are represented relative to CD45+ immune cells. Animals per treatment group n=3-5 from 2 independent experiments; Significance/p-values were calculated by one-way ANOVA and indicated as follows: *p < 0.05; **p < 0.01, ***p < 0.001; ns = not statistically significant.

**Supplementary Figure 7:**
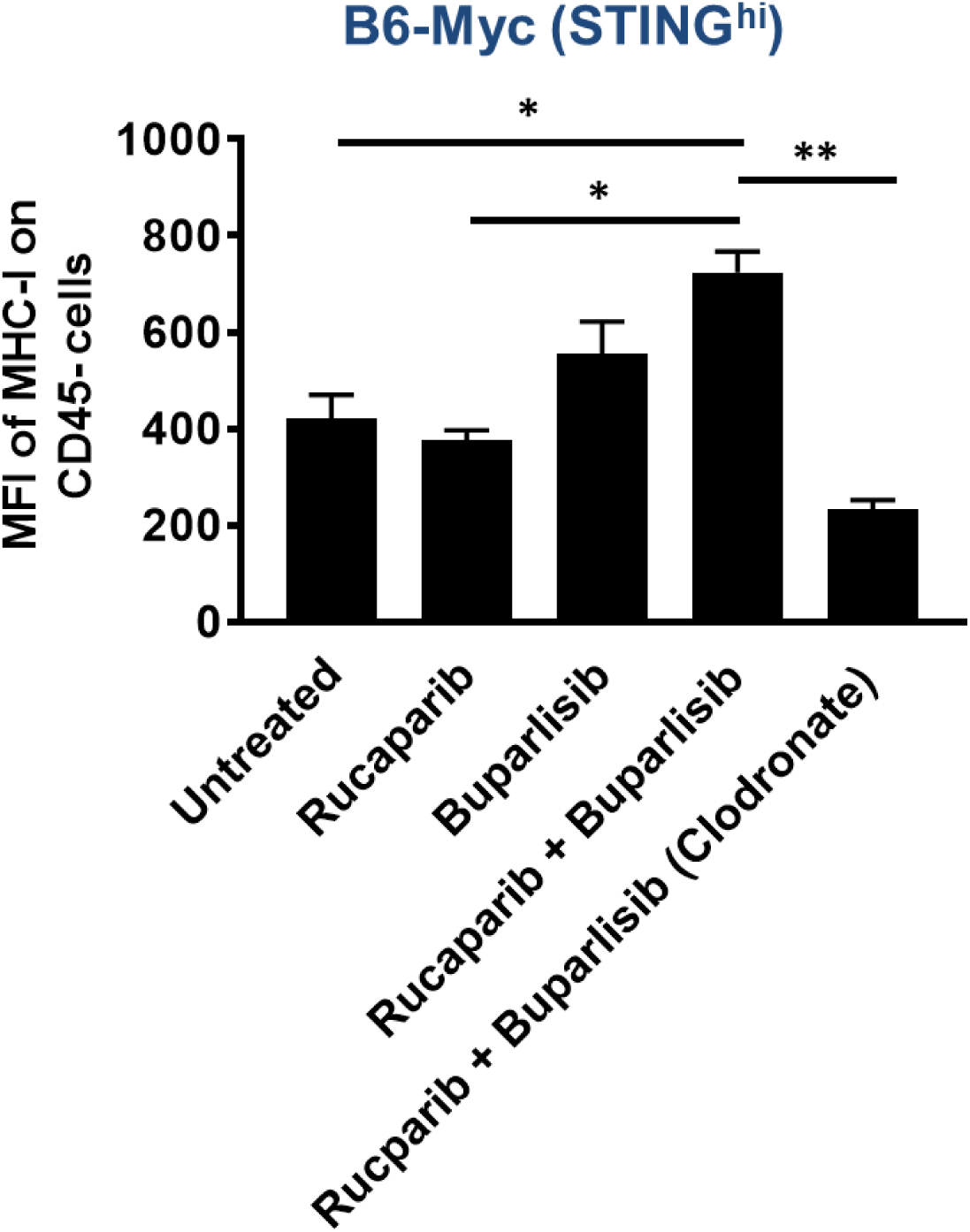
Concomitant clodronate treatment decreases MHC-I expression on CD45^-^ cells in B6-Myc (STING^hi^) tumor-bearing mice that were treated with PARPi/PI3Ki treatment. B6-Myc (STING^hi^) tumor-bearing syngeneic mice were treated with PARPi/PI3Ki +/- clodronate (to deplete macrophages) until untreated tumors reached 2500 mm^3^. Tumors were processed and stained for MHC-I by flow cytometry; n=3 animals per group; Significance/ p-values were calculated by one-way ANOVA, indicated as follows, **p < 0.01.

**Supplementary Figure 8:**
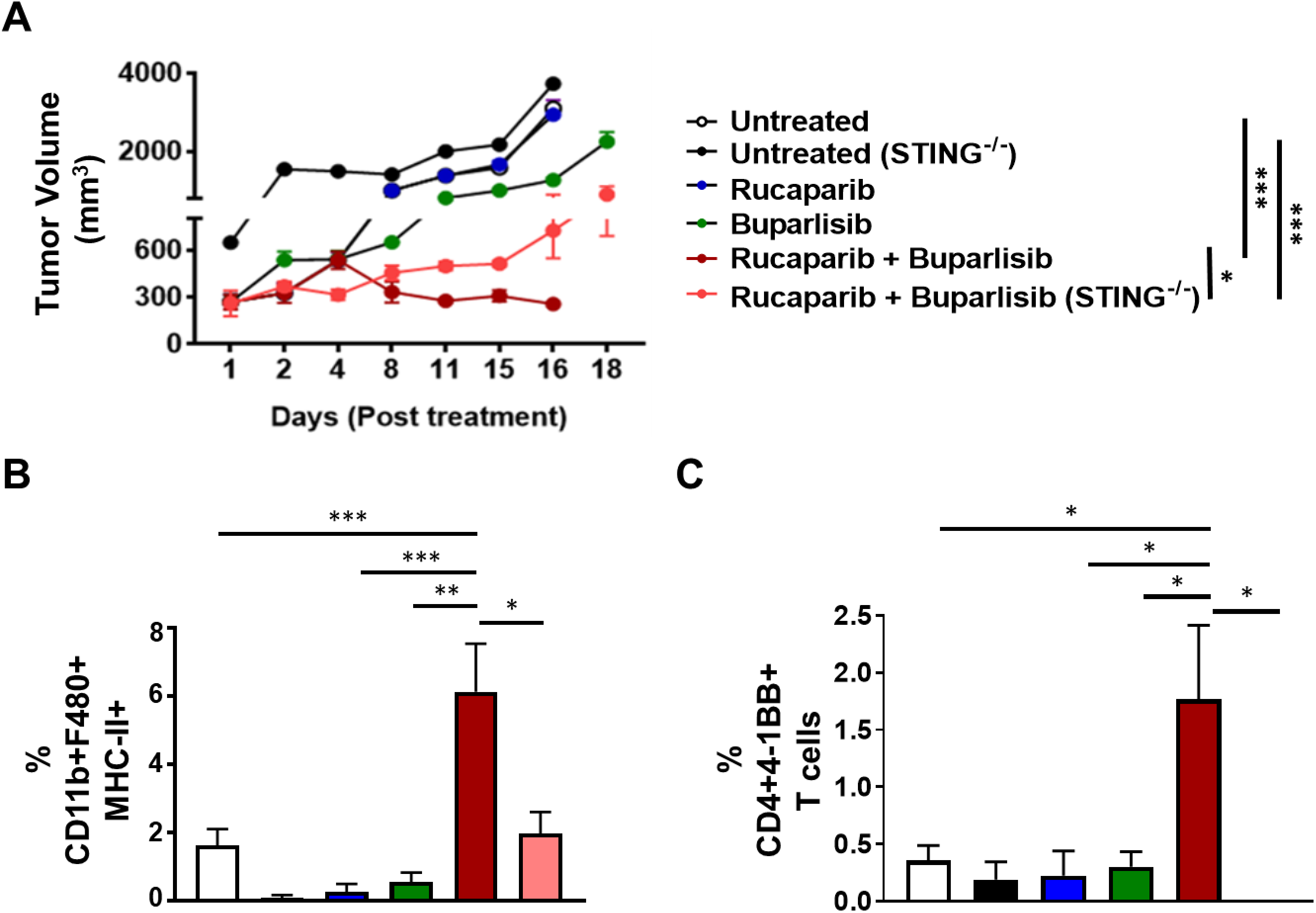
PARPi/PI3Ki combination therapy causes tumor regression in B6-Myc (STING^hi^) murine PC via host STING-dependent immune mechanism. C57Bl/6J^STING+/+^ and C57Bl/6J^STING-/-^mice were engrafted with B6-Myc (STING^hi^) tumor allografts and treated with the indicated drug(s) until untreated tumors reached approx. 2500 mm^3^. **(A)** Tumor volume curves for duration of treatment are shown. **(B)** Single-cell suspensions were generated from harvested tumors, and analyzed by flow cytometry, for activated macrophages **(B)** and activated T cells **(C)**. Data are represented relative to CD45+ immune cells. n=3 mice per group; Significance/ p-values were calculated by one-way ANOVA are indicated as follows, *p < 0.05; ****p < 0.0001.

**Supplementary Figure 9:**
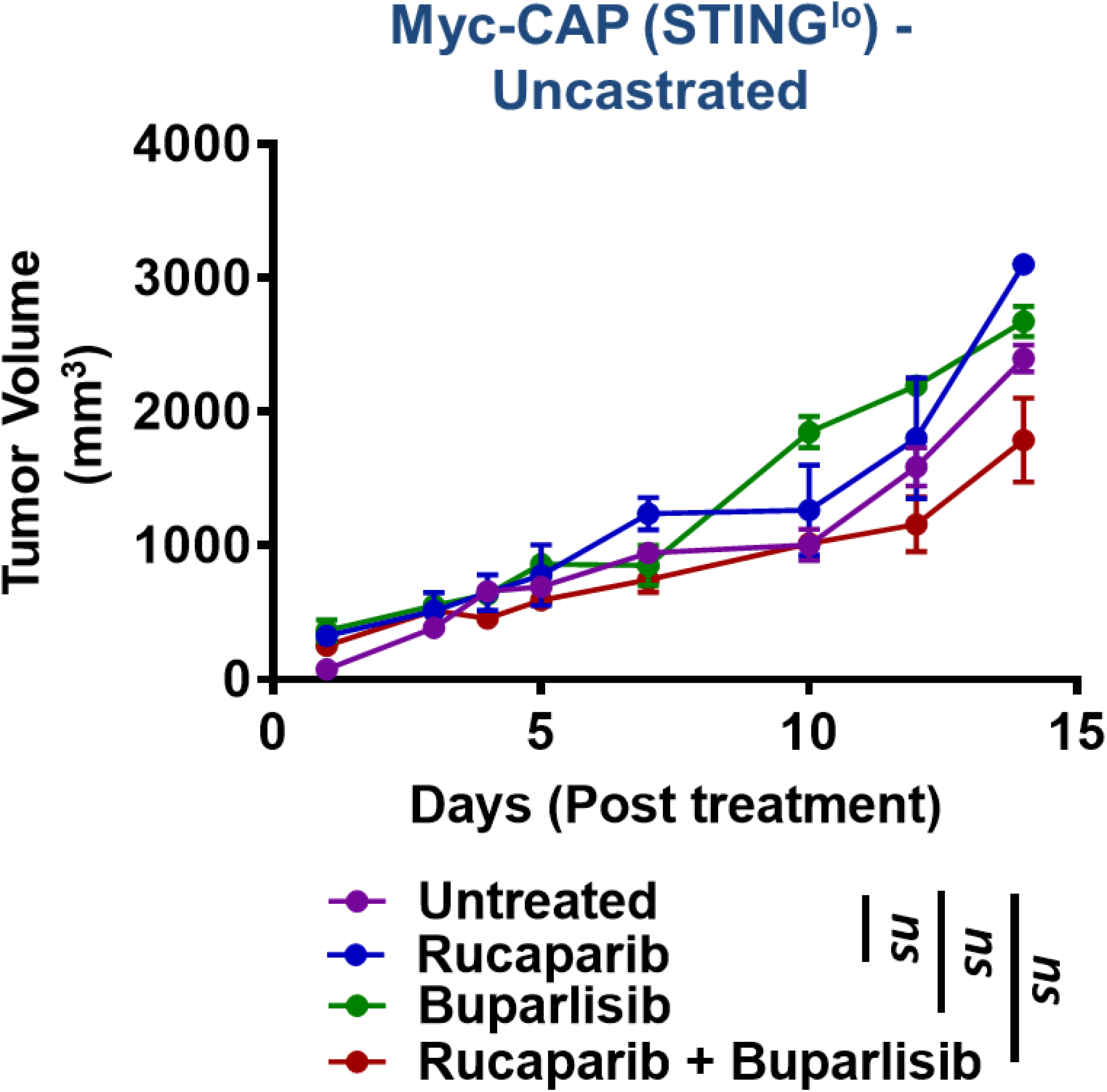
Myc-CAP (STING^lo^) tumors do not respond to PARPi/PI3Ki treatment in the absence of castration. Myc-CAP (STING^lo^) tumor-bearing syngeneic mice were treated with the indicated drug(s) until untreated tumors reached approx. 2500 mm^3^. Tumor volumes were recorded for duration of treatment; n=4 animals per group. Significance/p-values were calculated by one-way ANOVA and are indicated as follows: ns = not statistically significant.

**Supplementary Figure 10:**
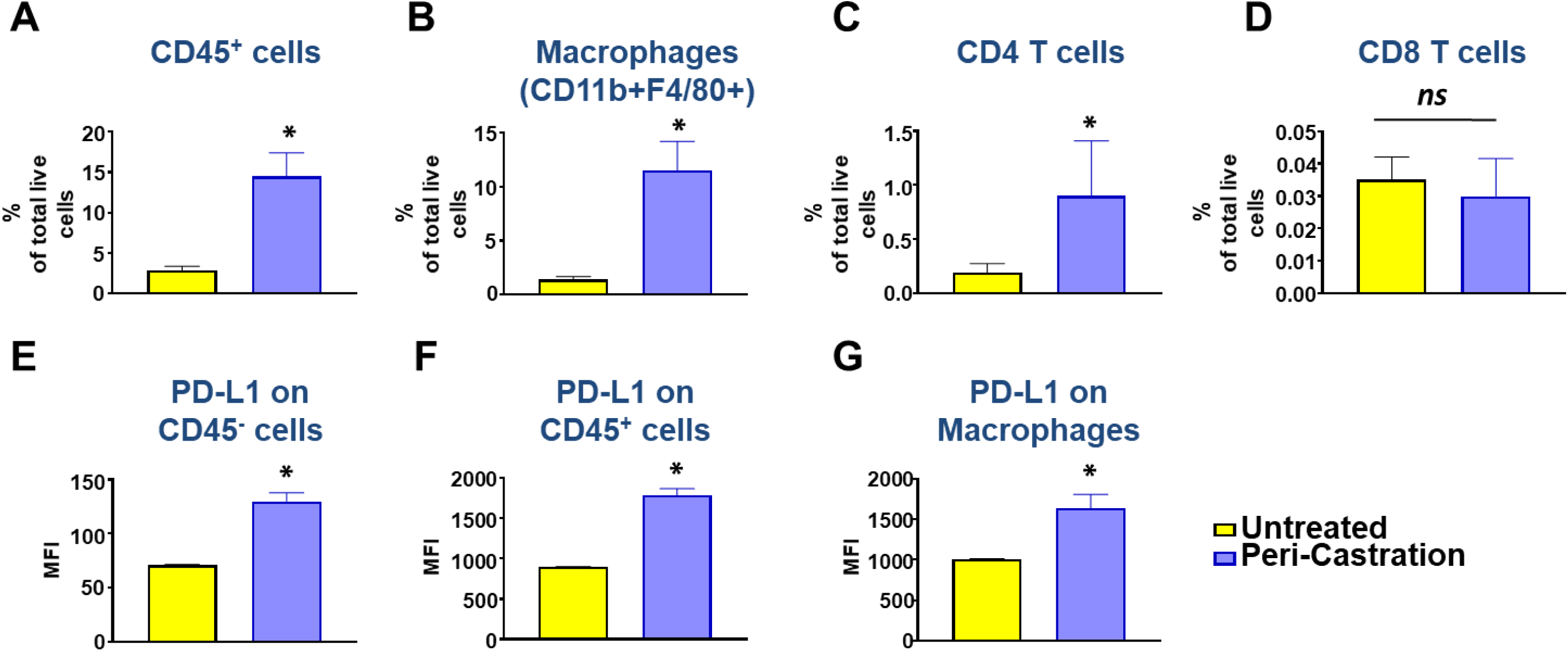
Chemical castration increases macrophage and CD4+ T cell infiltration and PD-L1 expression within the TME of Myc-CAP(STING^lo^) tumors. Myc-CAP (STING^lo^) tumor-bearing syngeneic mice were treated with degarelix (chemical castration) for 10 days. At the end of treatment, tumors were harvested and analyzed by flow cytometry for infiltration of CD45+ immune cells **(A)**, macrophages **(B)** and CD4/CD8 T cells **(C-D)**. Mean fluorescence intensity (MFI) are depicted for PD-L1 expression on CD45-/CD45+ cells **(E-F)** and TAMs **(G)** within the TME; n= 3-5 animals/group; Significance/ p-values were calculated by Un-paired t-test and indicated as follows *p < 0.05; ns= not statistically significant.

**Supplementary Figure 11:**
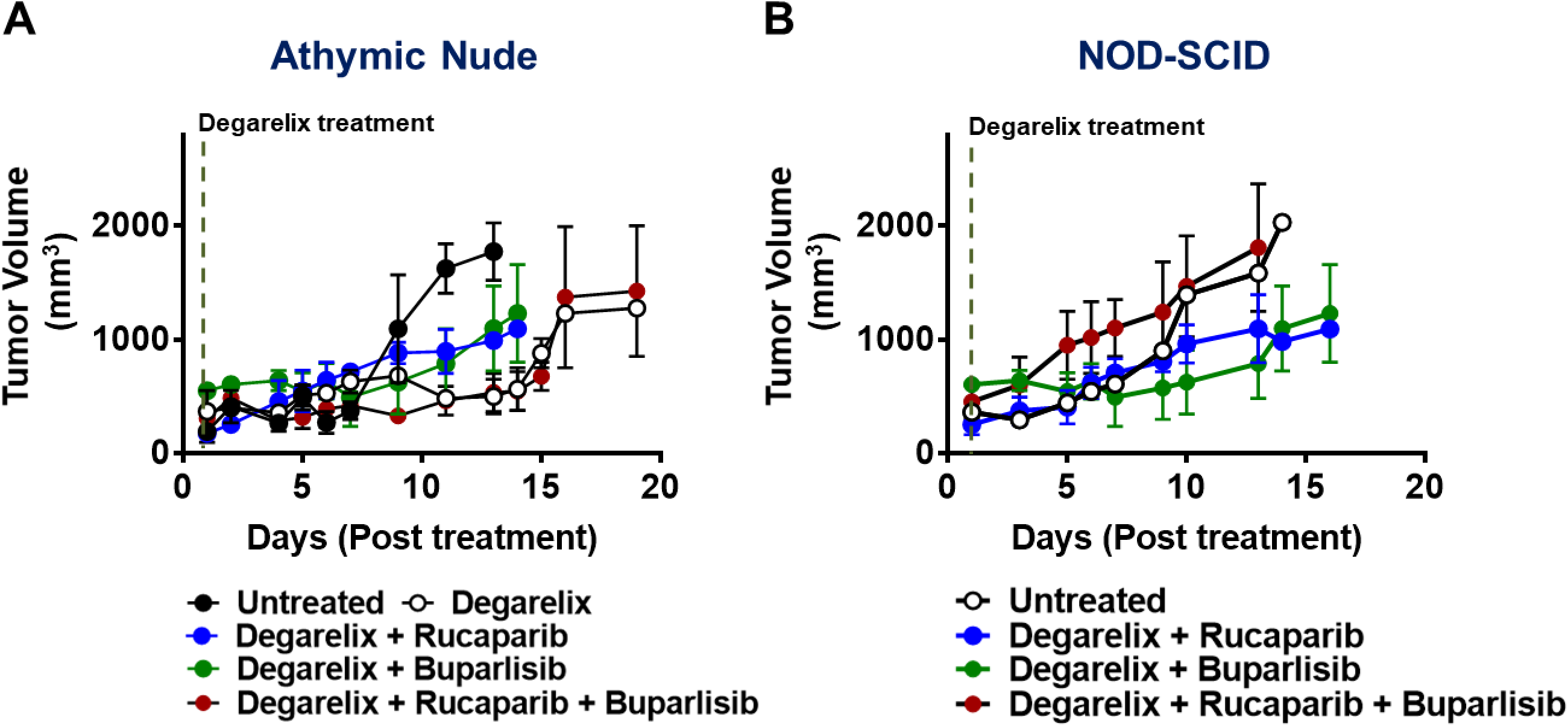
The anti-tumor response elicited by ADT/PARPi/PI3Ki is abrogated in Myc-CAP (STING^lo^)-tumor bearing athymic nude and NOD/SCID mice. Immunodeficient athymic nude **(A)** and NOD-SCID **(B)** mice were engrafted with Myc-CAP (STING^lo^) tumors and treated with the indicated drug(s) until untreated tumors reached approx. 2000mm^3^. Tumor volume curves for duration of treatment are shown. n=3-4 mice per group.

**Supplementary Figure 12:**
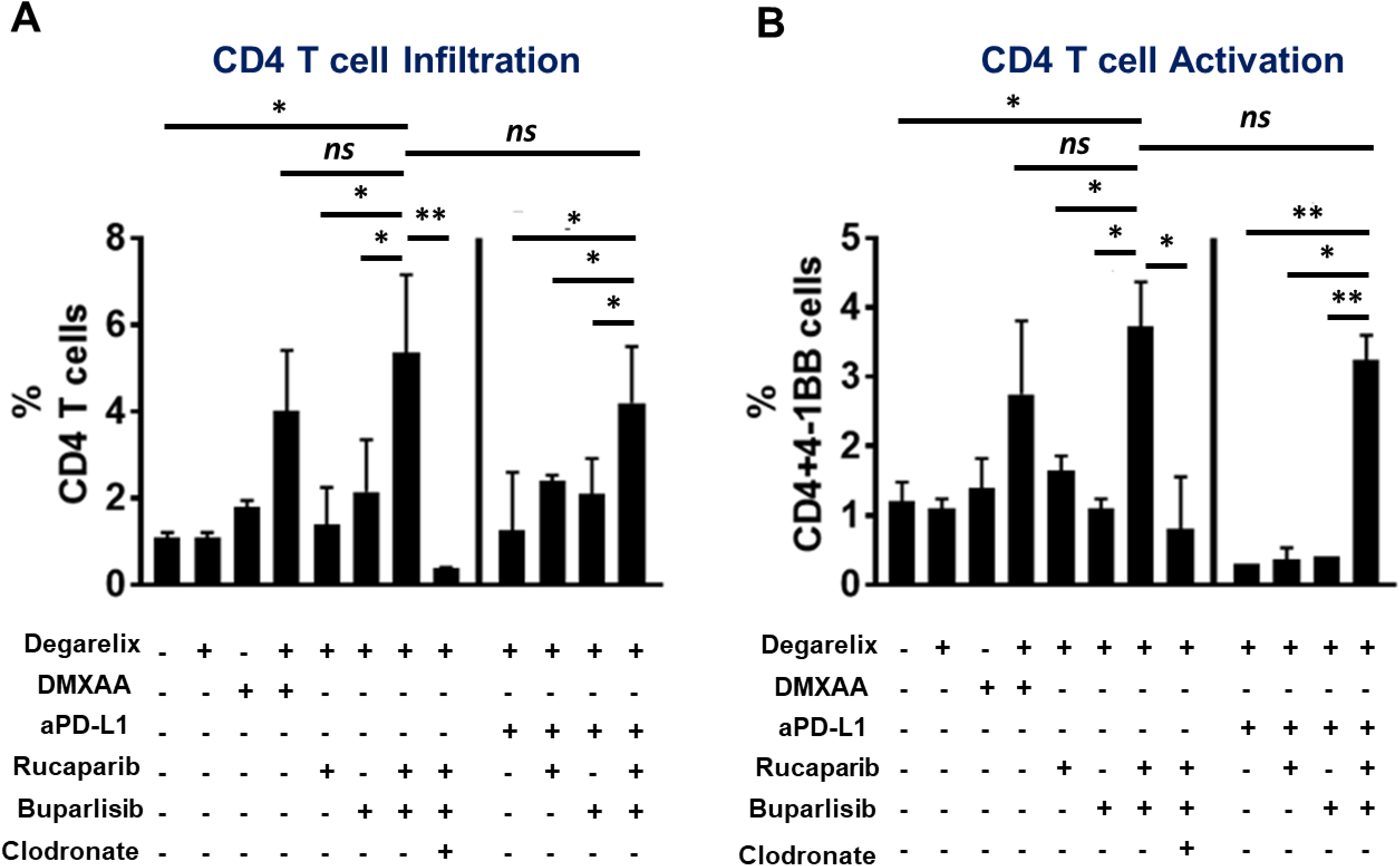
ADT/PARPi/PI3Ki with/without PD-L1 antibody treatment increases infiltration and activation of CD4 T cells within the TME of STING^lo^ Myc-CAP tumors. Myc-CAP (STING^lo^) tumor-bearing syngeneic mice were treated with the indicated drugs until untreated tumors reached approx. 2500 mm^3^. Single-cell suspensions were generated from harvested tumors and analyzed by flow cytometry for CD4 infiltration **(A)** and activation **(B)**. Data are represented relative to CD45+ immune cells. n=3-5 animals per treatment group from 2 independent experiments; Significance/ p-values were calculated by one-way ANOVA and are indicated as follows, *p < 0.05; **p < 0.01, ns = not statistically significant

**Supplementary Figure 13:**
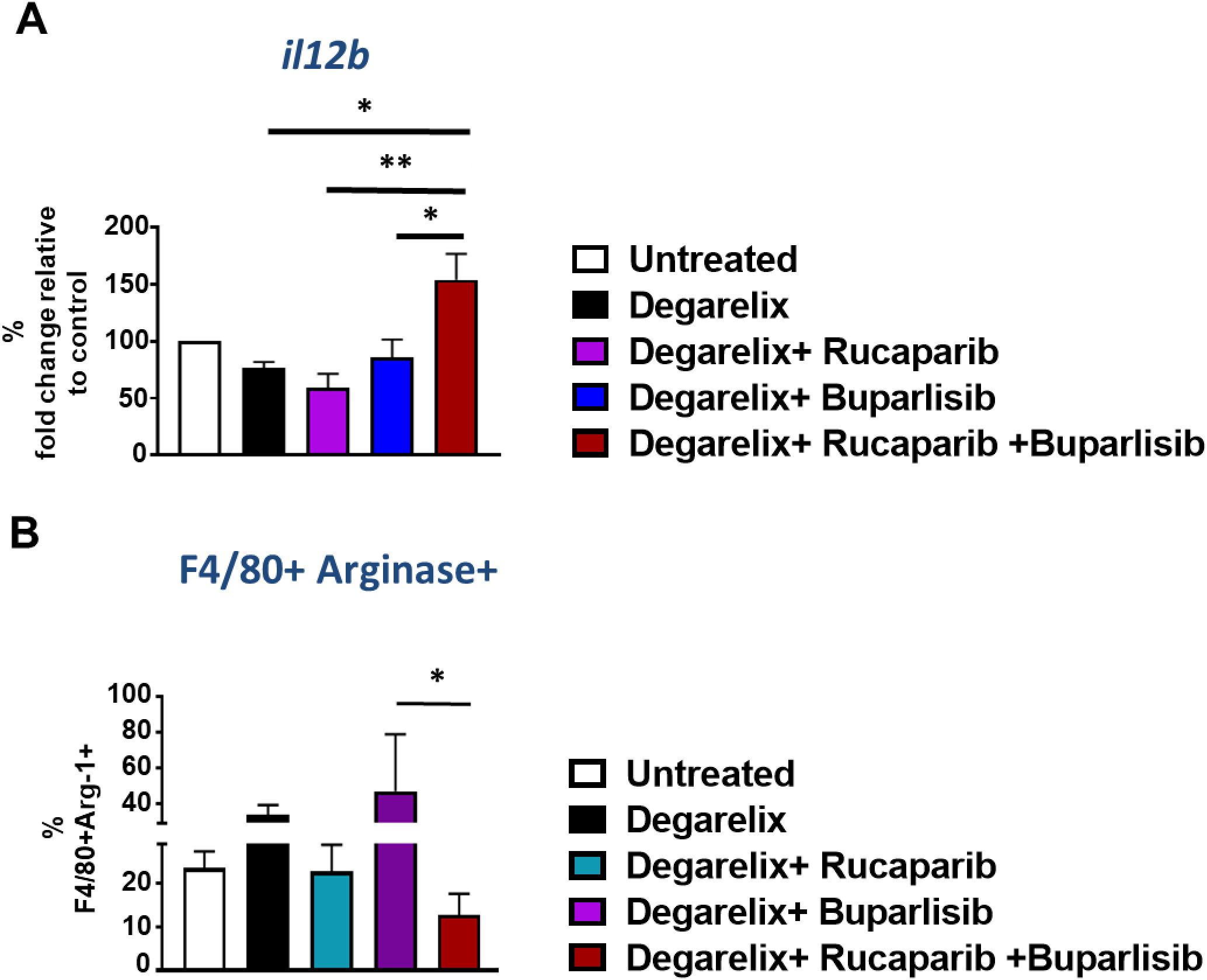
ADT/PARPi/PI3Ki treatment enhances M1 macrophage polarization within Myc-CAP (STING^lo^) tumors *in vivo*. **(A-B)** Myc-CAP (STING^lo^) tumor-bearing syngeneic mice were treated with the indicated drugs until untreated tumors reached approx. 2500 mm^3^. Tumors were harvested for qRT-PCR analysis to interrogate changes in *il12b* expression. **(B)** Single-cell suspensions were generated from harvested tumors and analyzed by flow cytometry for macrophage (F4/80+) Arginase I expression. n=3 animals/group; Significance/p-values were calculated by one-way ANOVA and are indicated as follows *p < 0.05, **p < 0.01.

**Supplementary Figure 14:**
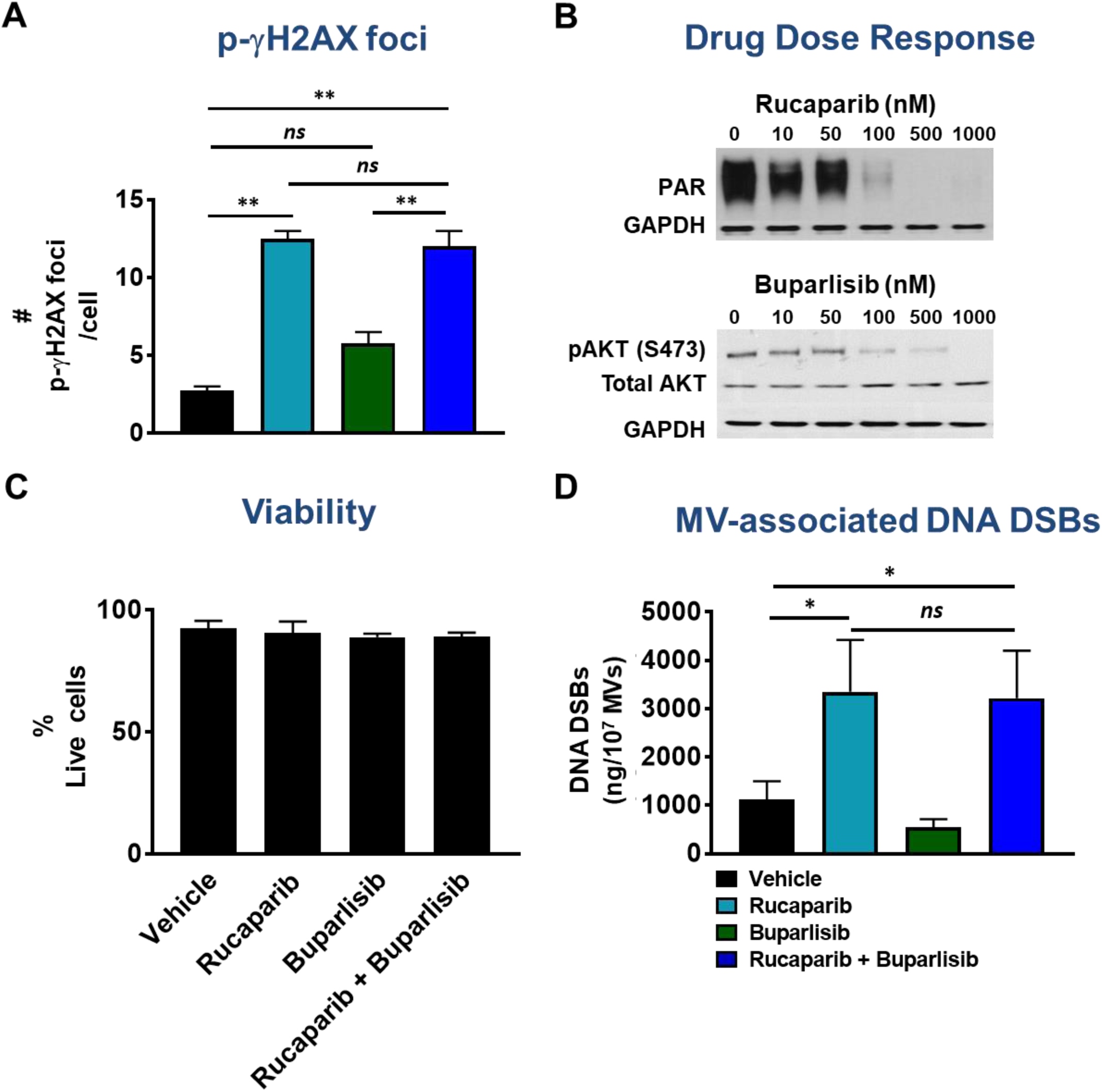
PARPi alone induces DNA DSBs, but no apoptosis even in combination with PI3Ki in Myc-CAP (STING^lo^) cancer cells. Myc-CAP cancer cells were treated with indicated drugs singly or in combination for 36 hours. **(A)** Cells were stained with anti-mouse specific p-γH2AX antibody and fluorescently labeled secondary antibody for determination of DNA DSBs, which were quantified by confocal microscopy. **(B)** Protein extracts from cells in **(A)** were analyzed for the indicated pharmacodynamic biomarkers by western blotting; **(C)** Annexin V-PI staining was done to assess frequency of live cells (Annexin V^-^ PI^-^) following drug treatment. **(D)** Ultracentrifugation was utilized to purify MVs from supernatants in treatment groups in **(A)** and associated DNA DSBs was quantified by Nanodrop. Independent experiments n=2. Significance/p-values were calculated by one-way ANOVA and are indicated as follows **p < 0.01; ***p < 0.001; ns = not statistically significant.

**Supplementary Figure 15:**
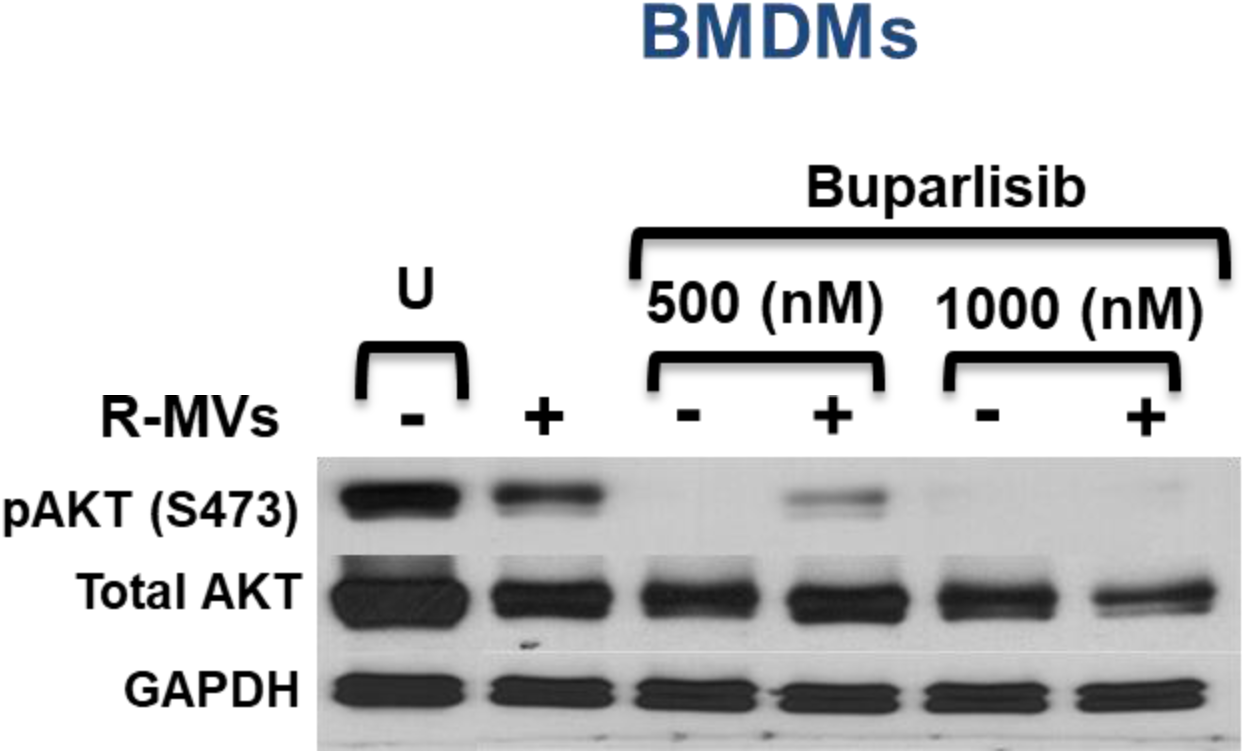
Buparlisib inhibits activation of the PI3K/AKT pathway within BMDMs. BMDMs were co-cultured with MVs isolated from Myc-CAP (STING^lo^) cancer cells following 36 hr treatment with rucaparib (R-MVs), in the presence or absence of buparlisib at the indicated concentrations for 24 hrs. Protein extracts were harvested for assessment of PI3K target inhibition by Western Blotting; U= Untreated.

**Supplementary Figure 16:**
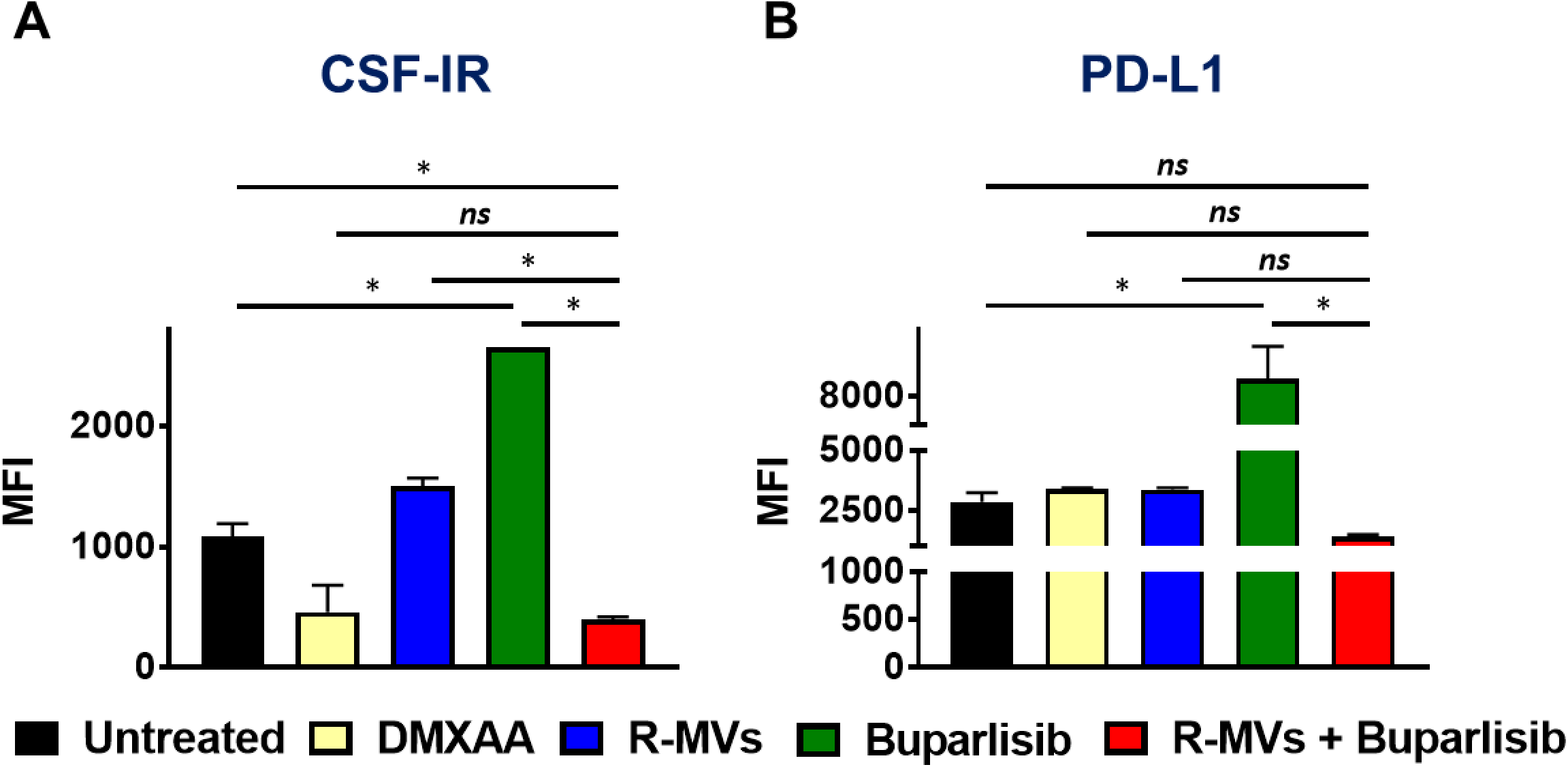
Conditioned medium from PARPi/PI3Ki treated Myc-CAP (STING^lo^) cells reprograms BMDMs from an M2 to M1 phenotype. BMDMs were co-cultured for 36 hours with supernatants from Myc-CAP (STING^lo^) cancer cells that were treated with the indicated drug(s) (rucaparib 0.5uM, buparlisib 1uM) for 36 hours. DMXAA (50ug/ml) was directly added to BMDMs. Following treatment, BMDMs were stained for expression of macrophage activation markers **(A)** MHC-II; **(B)** iNOS and suppressive marker **(C)** Arginase I on CD45+ CD11b+F4/80+ macrophages. **(D-E)** Supernatants were collected for determination of secreted chemokines by cytokine array; n=2 independent experiments; Significance/ p-values were determined by one-way ANOVA (A) / Un-paired t-test (B), *p < 0.05; **p < 0.01, ***p < 0.001, **** p< 0.0001, ns = not statistically significant.

**Supplementary Figure 17:**
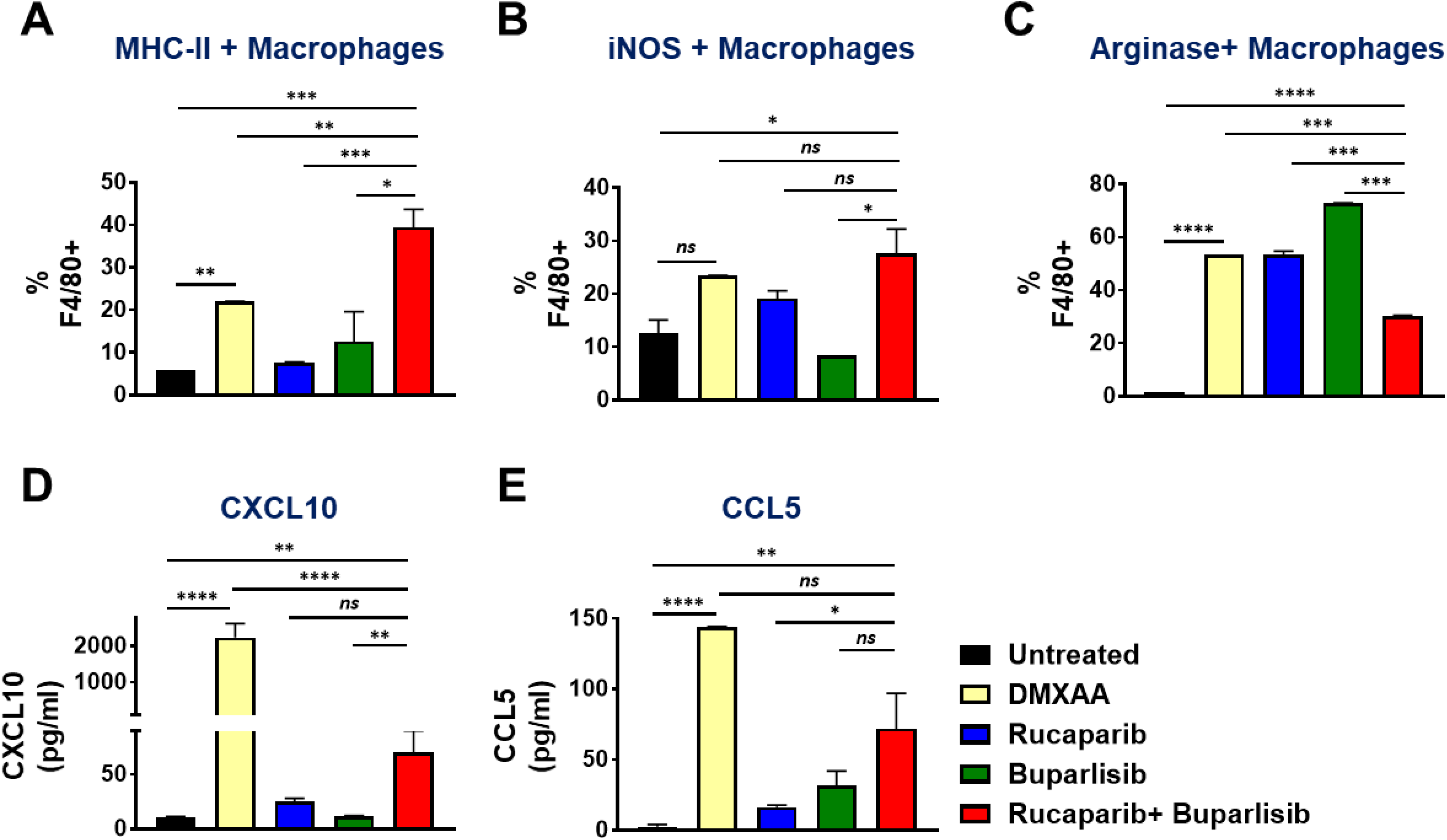
Conditioned medium from PARPi/PI3Ki treated Myc-CAP (STING^lo^) cells reprograms BMDMs from an M2 to M1 phenotype. BMDMs were co-cultured for 36 hours with supernatants from Myc-CAP (STING^lo^) cancer cells that were treated with the indicated drug(s) (rucaparib 0.5uM, buparlisib 1uM) for 36 hours. DMXAA (50ug/ml) was directly added to BMDMs. Following treatment, BMDMs were stained for expression of macrophage activation markers **(A)** MHC-II; **(B)** iNOS and suppressive marker **(C)** Arginase I on CD45+ CD11b+F4/80+ macrophages. **(D-E)** Supernatants were collected for determination of secreted chemokines by cytokine array; n=2 independent experiments; Significance/ p-values were determined by one-way ANOVA, *p < 0.05; **p < 0.01, ***p < 0.001, **** p< 0.0001, ns = not statistically significant.

## SUPPLEMENTARY TABLE LEGENDS

**Supplementary Table 1:** **Representative flow cytometric gating strategy, mean and standard deviations for specific immune subsets in human mCRPC and murine c-myc syngeneic tumor samples.** Single cell suspensions from human mCRPC biopsies, murine Myc-CAP and B6-Myc syngeneic tumors were stained with anti-human/mouse lineage-specific antibodies and analyzed by flow cytometry, as described in Figure 1. n=4 for human and n= 3-4 for murine samples. DC=dendritic cells; MDSC= myeloid derived suppressor cell, h= human, m= mouse.

| mCRPC patients | Gating strategy | Average (%) ± SD (%) |
| --- | --- | --- |
| Immune cells | Live+ CD45+ | 13 ± 1.43 |
| Non -Immune cells | Live+ CD45- | 87 ± 1.4 |
| Lymphoid | Live+ CD45+ CD3/CD45+ CD19+ | 22 ± 12 |
| Myeloid | Live+ CD45+ CD11b+ CD3- CD19- | 78 ± 8 |
| T cells | Live+ CD45+ CD3+ | 22 ± 12 |
| B cells | Live+ CD45+ CD3- CD19+ | 0 ± 0 |
| DCs | Live+ CD45+ CD11b+ CD11c+ | 6.4 ± 12 |
| Gr-MDSC | Live+ CD45+ CD11b+ HLA-DR- CD15<sup>hi</sup> CD33<sup>lo</sup> | 11.2 ± 3.7 |
| Mo-MDSC | Live+ CD45+ CD11b+ HLA-DR- CD15<sup>lo</sup> CD33<sup>hi</sup> | 13.6 ± 7.2 |
| Macrophage | Live+ CD45+ CD11b+ CD163+ CD68+ | 46.8 ± 9.4 |
| Activated Macrophages | Live+ CD45+ CD11b+ CD163+ CD68+ HLA-DR+ | 20.6 ± 10.6 |
| Un-activated Macrophages | Live+ CD45+ CD11b+ CD163+ CD68+ HLA-DR- | 79.4 ± 12.9 |
| Myc-CAP (STING<sup>lo</sup>) | Gating strategy | Average (%) ± SD (%) |
| Immune cells | Live+ CD45+ | 2.6 ± 0.9 |
| Non -Immune cells | Live+ CD45- | 97.4 ± 0.9 |
| Lymphoid | Live+ CD45+ CD3+ /CD45+ CD19+ | 22 ± 4 |
| Myeloid | Live+ CD45+ CD11b+ CD3- CD19- | 78 ± 4 |
| T cells | Live+ CD45+ TCRb+ | 20 ± 8 |
| B cells | Live+ CD45+ TCRb- CD19+ | 1.6 ± 0.9 |
| DCs | Live+ CD45+ CD11b+ CD11c+ | 2.2 ± 1.5 |
| Gr-MDSC | Live+ CD45+ CD11b+ MHC-II- Ly6G<sup>hi</sup> Ly6C<sup>lo</sup> | 0 ± 0 |
| Mo-MDSC | Live+ CD45+ CD11b+ MHC-II- Ly6G<sup>lo</sup> Ly6C<sup>hi</sup> | 33 ± 6.1 |
| Macrophage | Live+ CD45+ CD11b+ F480+ | 43.2 ± 7.2 |
| Activation state of Macrophages | Live+ CD45+ CD11b+ F480+ MHC-II+ | 35 ± 3 |
| Un-activated Macrophages | Live+ CD45+ CD11b+ F480+ MHC-II- | 65 ± 3 |
| B6-Myc (STING<sup>hi</sup>) | Gating strategy | Average (%) ± SD (%) |
| Immune cells | Live+ CD45+ | 26 ± 5.8 |
| Non -Immune cells | Live+ CD45- | 74 ± 5.8 |
| Lymphoid | Live+ CD45+ CD3+ /CD45+ CD19+ | 15.9 ± 2 |
| Myeloid | Live+ CD45+ CD11b+ CD3- CD19- | 84.1 ± 2.9 |
| T cells | Live+ CD45+ TCRb+ | 5.2 ± 2 |
| B cells | Live+ CD45+ TCRb- CD19+ | 10.2 ± 2 |
| DCs | Live+ CD45+ CD11b+ CD11c+ | 13.6 ± 3.5 |
| Gr-MDSC | Live+ CD45+ CD11b+ MHC-II- Ly6G<sup>hi</sup> Ly6C<sup>lo</sup> | 29.2 ± 4 |
| Mo-MDSC | Live+ CD45+ CD11b+ MHC-II- Ly6G<sup>lo</sup> Ly6C<sup>hi</sup> | 2.8 ± 1.1 |
| Macrophage | Live+ CD45+ CD11b+ F480+ | 39 ± 3 |
| Activation state of Macrophages | Live+ CD45+ CD11b+ F480+ MHC-II+ | 30.3 ± 3 |
| Un-activated Macrophages | Live+ CD45+ CD11b+ F480+ MHC-II- | 69.7 ± 3 |

